# The macronuclear genomic landscape within *Tetrahymena thermophila*

**DOI:** 10.1101/2023.10.02.560512

**Authors:** Romain Derelle, Rik Verdonck, Staffan Jacob, Michèle Huet, Ildem Akerman, Hervé Philippe, Delphine Legrand

**Author notes:** equal contribution (Hervé Philippe, Delphine Legrand).

## Abstract

The extent of intraspecific genomic variation is key to understanding species evolutionary history, including recent adaptive shifts. Intraspecific genomic variation remains poorly explored in eukaryotic microorganisms, especially in the nuclear dimorphic ciliates, despite their fundamental role as laboratory model systems and their ecological importance in many ecosystems. We sequenced the macronuclear genome of 22 laboratory strains of the oligohymenophoran *Tetrahymena thermophila*, a model species in both cellular biology and evolutionary ecology. We explored polymorphisms at the junctions of programmed eliminated sequences, and reveal their utility to barcode very closely related cells. As for other species of the genus *Tetrahymena*, we confirm micronuclear centromeres as gene diversification centres in *T. thermophila*, but also reveal a two-speed evolution in these regions. In the rest of the genome, we highlight recent diversification of genes encoding for extracellular proteins and cell adhesion. We discuss all these findings in relation with ciliate’s ecology and cellular characteristics.

**Impact Statement:** This is the first study of population genomics in the ciliate *Tetrahymena thermophila*. This bacterivore species plays an important role in aquatic trophic chains and is widely used as a model in cell and molecular biology, ecology, evolution or toxicology. As all ciliates, it contains a germline micronucleus and a somatic macronucleus. Sequencing of the macronucleus reveals that the centromeric region of the micronucleus are simultaneously a region of new gene diversification, as observed in other *Tetrahymena* species, and a region containing highly conserved genes. The results also confirm that the formation of the macronucleus from the micronucleus is highly imprecise. Interestingly, this process generates a genomic barcode that can discriminate cells derived from a given sexual reproduction event, allowing to study more finely population dynamics/history in nature.

**Data summary:** All data are fully provided in Supplementary Materials. The raw data of the 22 *Tetrahymena* genomes have been deposited in the Sequence Read Archive (https://www-ncbi-nlm-nih-gov.inee.bib.cnrs.fr/bioproject/PRJNA1012331). Accession numbers are listed in Table S1 (available in the online version of this article).

## Introduction

Intraspecific diversity is at the nexus of ecological and evolutionary dynamics [1, 2]. Its study can be done at several levels of biological organization, from molecules to populations. Among them, the genomic level has received much attention. Intraspecific genomic datasets allow to determine if genome diversity is a good proxy for phenome diversity (*i.e.*, the diversity of phenotypes), if evolutionary patterns detected at the intraspecific levels match those described at the interspecific level (*e.g.*, hot-spots of mutations, gene diversifications, genome size variations), or if the reconstruction of demographic, selective or ancestry patterns are consistent across the whole genome. Besides these evolutionary considerations, associations between reliable genome assemblies/annotations and phenome descriptions at the intraspecific level are fundamental in cellular biology and medicine to ascertain the functional role of specific genomic regions in a given biological process or disease [3]. Originally restricted to a few model organisms with often rather simple genomes, such eco-evolutionary and mechanistic aspects can now be investigated at the biodiversity level thanks to the recent advances in sequencing technologies.

*Tetrahymena thermophila*, a free-living ciliate, is a well-established model in cellular and molecular biology, which has led to major discoveries including identifications of dynein, telomere structure, telomerase and the first histone-modifying enzyme [4, 5]. It has greatly contributed to a better understanding of important genetic processes such as RNA interference and epigenetic regulation [6], and of the dynamics and regulation of cell cortical organization [7] and amitotic cell divisions [8]. In recent years, due to its easy maintenance and manipulation in laboratory microcosms [9, 10] and to its remarkable phenotypic attributes (*e.g.*, inducible caudal cilium [11, 12]), *T. thermophila* emerged as an experimental model in ecological and evolutionary studies. It contributed to a better understanding of the evolution of dispersal [13], and its interplay with adaptation and cooperation [14–16], and of phenotypic plasticity [17]. *T. thermophila* is an important actor of aquatic trophic chains, and thus serves as a model to study interspecific interactions, notably with its bacterial preys [18], and phages [19]. It is also widely used as an ecotoxicological model [20] and has been proposed as a bioindicator of polluted environments [21].

Like other ciliates, *T. thermophila* possesses two nuclear genomes with different roles during its lifecycle: the ‘germline’ micronuclear (MIC) and the ‘somatic’ macronuclear (MAC) genomes. The *T. thermophila* MIC genome is diploid, composed of five metacentric chromosomes totalling ∼157 Mb [22]. It is transcriptionally silent during vegetative growth and its role is mainly restricted to sexual life stages [5]. The polyploid (∼45n) MAC genome is composed of 181 chromosomes lacking centromeres. It totals ∼104 Mb, of which 48% encodes genes [23, 24]. It is transcriptionally active, controlling somatic cell functions during vegetative growth.

While the MIC conforms to the general rules of sexuality, the MAC formation is a ciliate-specific mechanism. Following each sexual reproduction, the meiotic products of the two progenitor cells fuse to give rise to a new nucleus that divides and generates the new MIC and MAC, which is a MIC-derived product. MAC formation is a sequential process involving the breakage of a duplicated version of the five MIC chromosomes into ∼225 MAC chromosomes in *T. thermophila* (of which 181 are ultimately conserved) on the basis of a 15-bp motif called Chromosome Breakage Sequence (CBS). Meanwhile, some repetitive genomic portions called Internal Eliminated Sequences (IES), mostly derived from transposable elements, are excised in a more or less precise process [25], to generate MAC Destined Sequences (MDS) that are finally reshuffled to form the final MAC genome. During MAC development, several endo-replication steps occur leading to a ∼45 ploidy level in *T. thermophila* [5], except for the small chromosome (20kb) encoding the 35S rRNA precursor gene that gets duplicated to ∼9,000 copies [26]. After the formation of the new MAC, the old MAC degenerates.

Variable numbers of chromosomes and ploidy levels are found across ciliates, with chromosome copy numbers up to ∼16 000 in the spirotrichean *Oxytricha trifallax*. In all cases, clonal divisions of the MAC are amitotic. This means that unequal copy numbers are propagated in daughter cells, theoretically leading to unpredictable drift of (i) chromosome copy numbers across asexual generations (but some regulatory mechanisms might exist [27–29]); and (ii) allelic composition, the so-called ‘phenotypic assortment’ [5]. In the case of *T. thermophila* (∼45n), the latter implies that, in the absence of selection, each cell lineage fixes 99% of its loci in ∼200 asexual generations. Amitosis offers an unusual panel of somatic genomic variants on which selection can act at each asexual generation [30]. For instance, in the presence of toxic cadmium ions, both chromosome copy number variation and paralogous expansion of metallothionein have been observed in *T. thermophila* in a few asexual generations [31]. The evolutionary origin of this somatic genome remains unclear but a role in the domestication of transposable elements [32–34], as well as a role in phenotypic plasticity through somatic selection [30], have been proposed.

Several ciliate MAC reference genomes have been published, including of species of the genera *Tetrahymena* [35], *Paramecium* [29, 36], *Oxytricha* [37] or the heterotrichean *Stentor* [38]. However, as for most eukaryotic microorganisms, very few intraspecific datasets are available, except in the *Paramecium* genus [29, 36]. To the best of our knowledge, comparative genomic studies in the genus *Tetrahymena* have only been performed at the inter-specific level [35]. The mapping of recently evolved genes on the MIC assembly of *T. thermophila* revealed their concentration on pericentromeric and subtelomeric chromosome regions, suggesting these regions as genome innovation centres [35]. While this dataset sheds light on intra-genus genome evolution, the extent and organization of genomic diversity at the intra-specific scale is warranted.

Here, we analyse the polymorphism of *T. thermophila* MAC genome after sequencing 22 laboratory strains originating from strains collected by Professor Paul Doerder between 2002 and 2009. Since their sampling, these strains have been maintained under laboratory conditions, and have been used in experimental microcosms (*e.g.*, [11, 15]). They differ in many phenotypic features, including gross morphology, mobility, thermal optimum, growth parameters and dispersal ability (*e.g.*, [11, 39, 40]). Our aim is to provide a new MAC genomic resource for the model *T. thermophila* by generating the densest intraspecific sampling to date. The MAC polymorphism landscape of *T. thermophila* includes two types of Single-Nucleotide Polymorphisms (SNPs; Figure 1), all subject to phenotypic assortment: (i) SNPs originating from the two MIC chromosomes and retained by phenotypic assortment, or ‘MIC-derived-SNPs’, that should be the most frequent, and (ii) SNPs resulting from *de novo* mutations occurring any time during the asexual phase, or ‘MAC-*de novo*-SNPs’, that should be much less frequent (but depending on the length of asexual phase). In this paper, we describe diversity metrics at the genome scale using both MIC-derived-SNPs and fixed or high-frequency MAC-*de novo*-SNPs (see Methods). We also describe the variability at MDS junctions created by the imprecise process of IES excision occurring during the MAC formation of *T. thermophila*. We then explore the selective forces acting especially on coding regions. Finally, we compare these genomic characteristics observed at the within-species level to the evolutionary patterns reported at the interspecific level between *Tetrahymena* species [35].

**Figure 1:**
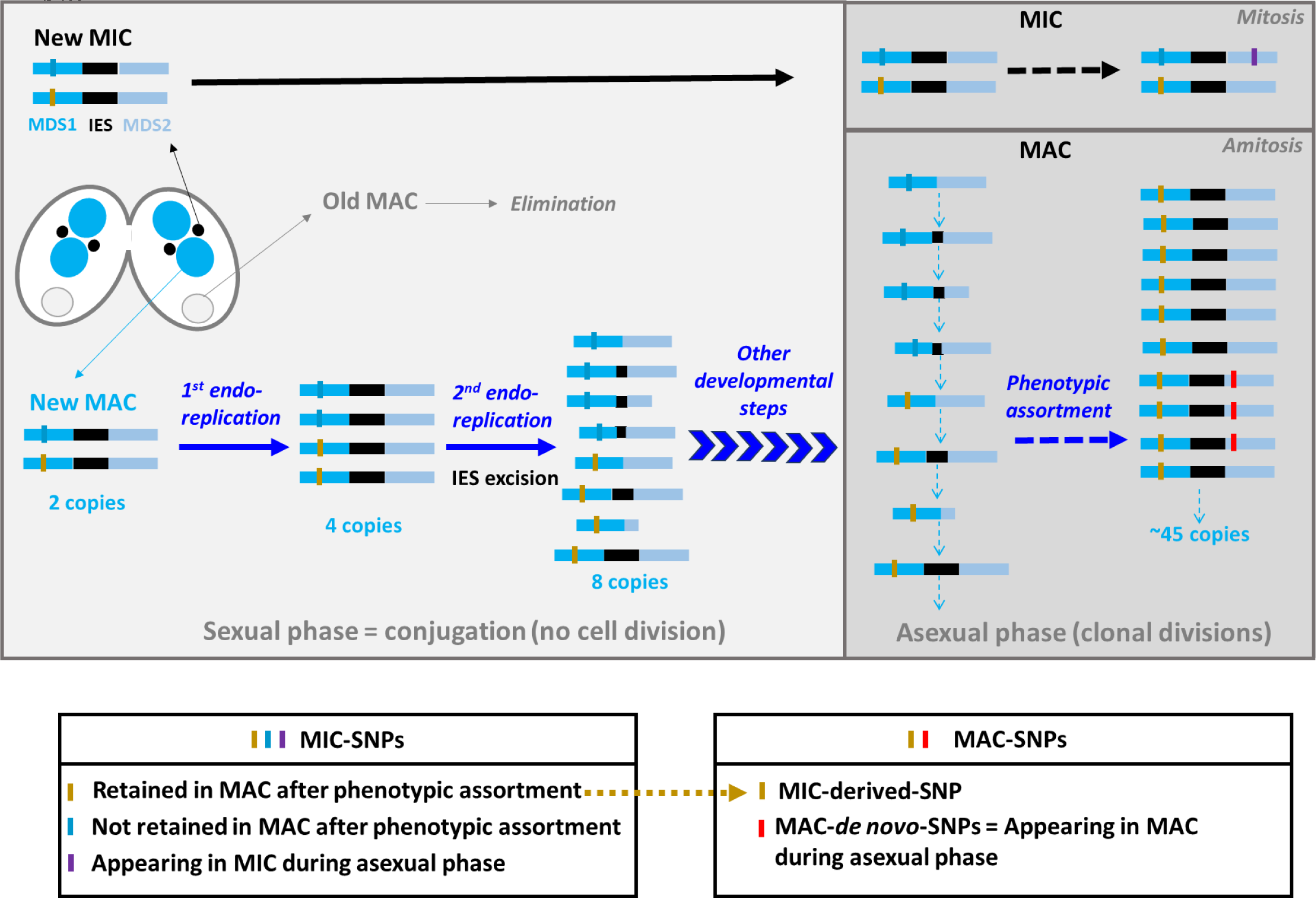
The MAC polymorphism landscape of *T. thermophila*. After exchange and fusion of the two parental meiotic products, two mitoses of the newly formed MIC in each conjugant cell (represented on the left) give rise to four nuclei, two staying MICs (in black, one is later eliminated), the two others being processed as new MACs (in blue), while parental MACs lately degenerate (in grey) (see the complete conjugation process in Figure 10 of [75]). MAC’s development relies on intensive genome remodelling giving rise to a mosaic of SNPs with different origins. For simplicity, we illustrate here a unique MIC locus composed of two MAC Destined Sequences (MDS1 and MDS2, horizontal blue bars) separated by one Internal Eliminated Sequences (IES, horizontal black bar). This locus is composed of two parental alleles characterized by a point mutation (blue and yellow vertical bars = MIC-SNPs). A first endoreplication round in young MACs increases the genomic copy number to four, without remodelling. During the second round of endoreplication, IES excision, a highly imprecise process in *T. thermophila* can give rise to up to eight variants. From the top to the bottom for copies with the blue allele: complete excision of the IES, small incomplete IES excision, small incomplete IES excision plus excision of a large part of MDS2, small incomplete IES excision plus excision of a small part of MDS1. From the top to the bottom for copies with the yellow allele: complete excision of the IES, large incomplete IES excision, complete IES excision plus excision of a large part of MDS2, retention of the full IES. These variants are then propagated throughout the MAC development, which continues after conjugants’ separation, *i.e.*, during the juvenile phase of the vegetative cycle. The whole process of MAC development notably includes the breakage of the five MIC chromosomes into smaller chromosomes and the stabilization of the MAC content to ∼45 copies (except the chromosome encoding the 35S rRNA). During the asexual phase, diploid MICs propagate mitotically, while MACs propagate amitotically. MAC genomes are therefore subject to phenotypic assortment, which leads to the random fixation (or loss) of SNPs derived from the MICs (yellow bar in the right “MAC” panel = MIC-derived-SNPs). Phenotypic assortment also retains one variant among those potentially created by imprecise excision of IES. Throughout the asexual phase, *de novo* MAC mutations (red vertical bars = MAC-*de novo*-SNPs) will segregate as well according to the phenotypic assortment process. The purple vertical bar represents a *de novo* MIC-SNP that can only be transmitted to a MAC after another conjugation event.

## Methods

### Culture of *T. thermophila* strains

The 22 *T. thermophila* strains used in this study were kindly provided in 2014 by Nicolas Schtickzelle (Biodiversity Research Center of Louvain la Neuve, Belgium). They were initially collected in the field by Pr Paul Doerder (Cleveland State University) between 2002 and 2009 and then referenced in the *Tetrahymena* stock centre (Cornell University, USA, Supplementary Material 1). In the Schtickzelle lab, each strain was maintained in axenic liquid growth media (2% Difco proteose peptone, 0.2% yeast extract, 1X PPYE) at 27°C within 24-well culture plates by transferring every ∼7 days. In 2014, the strains were sent to the Station d’Ecologie Théorique et Expérimentale of Moulis (France) where they were maintained in similar culture plates but at 23°C in a less concentrated axenic media (0.6% Difco proteose peptone, 0.06% yeast extract, 0.3X PPYE) with transfer every ∼10 days.

### DNA extractions and sequencing

We cultured each strain in a higher volume than the one used for routine culture by placing 50µL of 7-day old cultures in 50mL Falcon tubes filled with 20 mL of 0.3X PPYE medium. After 12 days of growth at 23°C (mid/end-stationary phase), we concentrated the cells by centrifuging each tube at 5,000 rpm for three minutes and discarding the supernatant. We added 200 µL of PBS to each cell pool and performed the 22 DNA extractions using the Buccal Swab protocol of the ReliaPrep™ gDNA Tissue Miniprep System kit (Promega). We adjusted the lysis time to two hours before performing the RNAse treatment. We did not separate the MIC and MAC during the extraction, meaning that we expect the presence of both MIC (2n) and MAC (∼45n) sequences before bioinformatic filtering (see below). The quantity and quality of extracted DNA were assessed with a Dropsense (Trinean, Supplementary Material 1) and good quality of extractions was confirmed by the Novogene company (Supplementary Material 2), who performed the library preparation and sequencing. An Illumina HiSeq sequencer was used to generate 5.2 Gb raw data/sample of 150 bp paired-end sequences (50x expected coverage of the MAC genome) from DNA libraries obtained after a PCR-free protocol, which decreases both unequal genomic coverage and the probability of erroneous base calls due to technical biases.

### Evolutionary distances and phenetic trees

Reads of our 22 strains were trimmed using Trimmomatic v0.39 [41] with recommended settings for Illumina paired-end reads and assembled into genome assemblies by SPAdes v3.14.1 [42] using the options ‘--only-assembler --careful’. MAC and mitochondrial chromosomes were reconstructed by scaffolding the SPAdes contigs onto the reference SB210 genome using RagTag v1.1.0 (https://github.com/malonge/RagTag). Then, all-versus-all pairwise Jukes-Cantor distances were estimated among the 22 genome assemblies and the reference SB210 MAC genome using andi v0.23 [43]. The resulting distances matrix was used by Ninja v1.2.2 [44] to infer MAC and mitochondrial neighbour-joining trees.

### Adjustments of the reference SB210 MAC genome

For all analyses, we used the complete SB210 MAC genome assembly published in [24] after performing several modifications described in Supplementary Material 3. Briefly, we polished the assembly using Pilon v1.23 [45] and Illumina reads from [24], renamed the MAC chromosomes according to their position in the MIC genome and built a new gene prediction mainly based on the Ensembl gene prediction v2.2.48. This corrected assembly and gene prediction are available for download at https://zenodo.org/record/7991891. The slim Gene Ontology annotation was downloaded from Ensembl protists biomart (http://protists.ensembl.org/biomart/martview/). In total, our annotation comprised 26,018 protein-coding genes.

### Read processing and SNP calling

The trimmed reads were mapped onto the SB210 MAC and mitochondrial genomes using the ‘mem’ algorithm of Bwa v0.7.17 [46]. The mappings were filtered to retain those with a mapping score of at least 30 and used to infer coverages and to identify SNPs and indels using SAMtools v1.10, BCFtools v1.9 [47] and BEDTools v2.28 [48]. All command lines used in this study are available in Supplementary Material 3. SNPs and indels were then filtered using a custom python script to keep those supported by at least three reads. This filtering limited false positives due to sequencing errors, but more importantly to the presence of reads from the MIC genome with alleles that were not retained in MACs after phenotypic assortment (*i.e.*, MIC-SNPs only present in the MIC genome) and the rare *de novo* MIC-SNPs that accumulated in MICs after the formation of MACs (purple bar in Figure 1). These MIC-SNPs are present in 1n in DNA extracts, and should therefore be present at an expected coverage of ∼1x in our 50x dataset. We also only kept SNPs and indels supported by at least one forward and one reverse alignment to limit mapping errors. Then, we chose to only consider variant calling positions that had a coverage of at least 20x (to avoid repeat regions) and below 80x (to avoid segmental duplications not present in the reference genome). Our dataset should thus mainly be composed of MIC-derived-SNPs and high frequency/fixed MAC-*de novo*-SNPs (Figure 1). However, MIC-SNPs not retained in MACs after phenotypic assortment, as well as *de novo* MIC mutations, could still be present at mid-high frequency in this dataset due to potential hyperdiploidy of some MIC chromosome regions in some strains. Aneuploidy can indeed occur in *T. thermophila* MICs (between 1n and up to almost 6n [49] as a consequence of multiple vegetative reproduction cycles [50]). Hyperploidy should randomly affect MIC chromosome regions and strains, meaning that the origin of mid-high frequency mutations is difficult to ascertain when performing MAC sequencing. As this could influence estimations of MAC mutational and diversity parameters, we decided to only consider positions for which the nucleotide different from the reference SB210 genome had a minimum frequency of 30x when calculating nucleotide polymorphism metrics. In all 22 strains combined, they represented 95.6% of all varying positions. We thus believe that after this last filtering step, our dataset was a reliable representative of the MAC genomic landscape. For simplicity, positions presenting more than two character-states were discarded from all analyses.

### Nucleotide polymorphism metrics

Polymorphism metrics, *i.e.*, nucleotide diversity (calculated as in [51]), the number of non-synonymous over the number of synonymous substitutions pN/pS [52], and the Direction of Selection DoS [53] were calculated using custom python scripts. DoS measures the direction and extent of selection by comparing the number of non-silent (pN) and silent polymorphisms (ps) to the number of non-silent (dN) and silent (dS) substitutions per locus, avoiding statistical bias due to low sampling sizes sometimes observed with other selection metrics like the MacDonald and Kreitman test or the Neutrality Index [53]. DoS is positive in case of adaptive evolution, equals zero in case of neutral evolution, and is negative in case of slightly deleterious mutations segregating. For pN/pS calculations, we kept only one representative of the seven genomic clusters (see Results) and excluded the sequence of strains having a frameshift at any position of the gene (keeping the other strains). Then, the ratios were computed for genes having at least one synonymous and one non-synonymous SNP, resulting in 21,804 genes with pN/pS values among the 26,018 protein-coding genes. For the DoS, which is calculated as dN/(dN+dS) - pN/(pN+pS), we used pN and pS values mentioned above and the pairwise dN (Ka) and dS (Ks) values and orthologous groups published in [35]. Nineteen orthologous groups with a high number of *T. thermophila* genes were excluded (more than 15 *T. thermophila* genes, cut-off chosen to keep a reasonably low number of genes per orthologous group), and for the remaining 17,283 orthologous groups, the dN and dS values for each *T. thermophila* gene was calculated as the median of all pairwise values between this gene and the genes of the eight other available *Tetrahymena* species. The DoS values were finally calculated for all genes having orthologs in the *Tetrahymena* genus (available dN and dS) and dN+dS and pN+pS values different from 0, resulting in 16,892 *T. thermophila* genes with DoS values. We also computed pN/pS and DoS distributions of gene clustered by their annotation using the slim Gene Ontology (GO) [54] to determine which functional pathways evolved especially fast or slowly in the *T. thermophila* species.

To test for a functional enrichment among genes present around MIC centromeric regions, we created four gene datasets. The first one was composed of the MAC chromosomes corresponding to the MIC centromere position (MAC 24, 62, 96, 125 and 168). The three others were composed of these five MAC chromosomes plus one, two or three chromosomes around them. Gene counts in these regions can be found in Supplementary Material 4. For each of these datasets, we calculated the mean DoS, pN/pS and dN/dS and tested for functional enrichment using g:profiler [55] among the genes having higher values than the mean regional values (*i.e.*, genes evolving faster compared to the others present in the considered regions). Since we observed a significant decrease in the number of genes for which orthologs were found in the other species of the *Tetrahymena* genus around MIC centromeres (no available DoS estimation), we also tested for functional enrichment among genes that had pN/pS estimates but no DoS estimates. These genes should correspond either to recent (duplicated) genes originating after the *T. thermophila* divergence or to genes difficult to align between species of the *Tetrahymena* genus. Results were highly similar between the four datasets, and are thus hereafter presented as a whole. As a control to verify that the detected functional enrichments were specific to the MIC centromeric regions, we created a final dataset composed of the whole genome except MAC chromosomes hosting MIC centromeres plus the three chromosomes around and applied the same methodology.

## Results and Discussion

### Mapping, SNP calling and evolutionary distance

When mapped on the SB210 MAC reference genome (Supplementary Material 3), the 22 Illumina datasets had an average coverage between 47.6 and 61.7x (Supplementary Material 5A). The percentage of the SB210 MAC genome mapped at a minimum coverage of 20x varied among strains between 91.6 and 99.5%. Keeping variant calling positions supported by at least three reads and at least one forward and one reverse mapping, and considering only positions that had a coverage between 20x and 80x (see Methods), this percentage varied between 88.3 and 98.5%. After excluding positions with more than two character-states (no more than a few hundred per strain), a total of 2,326,587 biallelic positions were found, corresponding to 2.25% of the MAC genome. This value is similar to those obtained in the *Paramecium* genus [36]. Indeed, based on between 10 and 13 isolates resequenced with comparable coverage in five *Paramecium* species, 0.7% to 7.4% of the MAC genome was found polymorphic [36]. The number of differences compared to the SB210 reference genome varied greatly, from 48 in strain D18 to 1,129,251 in strain D12 (Supplementary Material 5B), representing ∼0 to ∼1.2% of divergence with the SB210 strain.

We estimated evolutionary distances between MACs of our 22 strains plus the reference SB210 by calculating Jukes-Cantor distances between all pairs of MAC genome assemblies. As shown by the Neighbour-Joining (NJ) tree inferred from these distances (Figure 2A), our dataset was composed of seven clusters of highly similar strains: D3/D6/D12, D10, D1, D11, D5, D16 and a large cluster comprising 14 strains nearly identical to the reference SB210 strain (referred hereafter as ‘SB210-like strains’). Assuming that MIC-derived-SNPs are the most frequent type of SNPs in the MAC genomic landscape of *T. thermophila*, these seven clusters are representative of seven distinct MICs. We also constructed a NJ tree including the MAC genomes of *T. malaccensis*, *T. elliotti* and *T. borealis* (Supplementary material 6), which are among the closest relatives of *T. thermophila* [35, 56]. The 22 strains formed a clade distant from other *Tetrahymena* species suggesting that, despite the large divergence observed between the seven clusters, all 22 strains belong to the same species.

**Figure 2:**
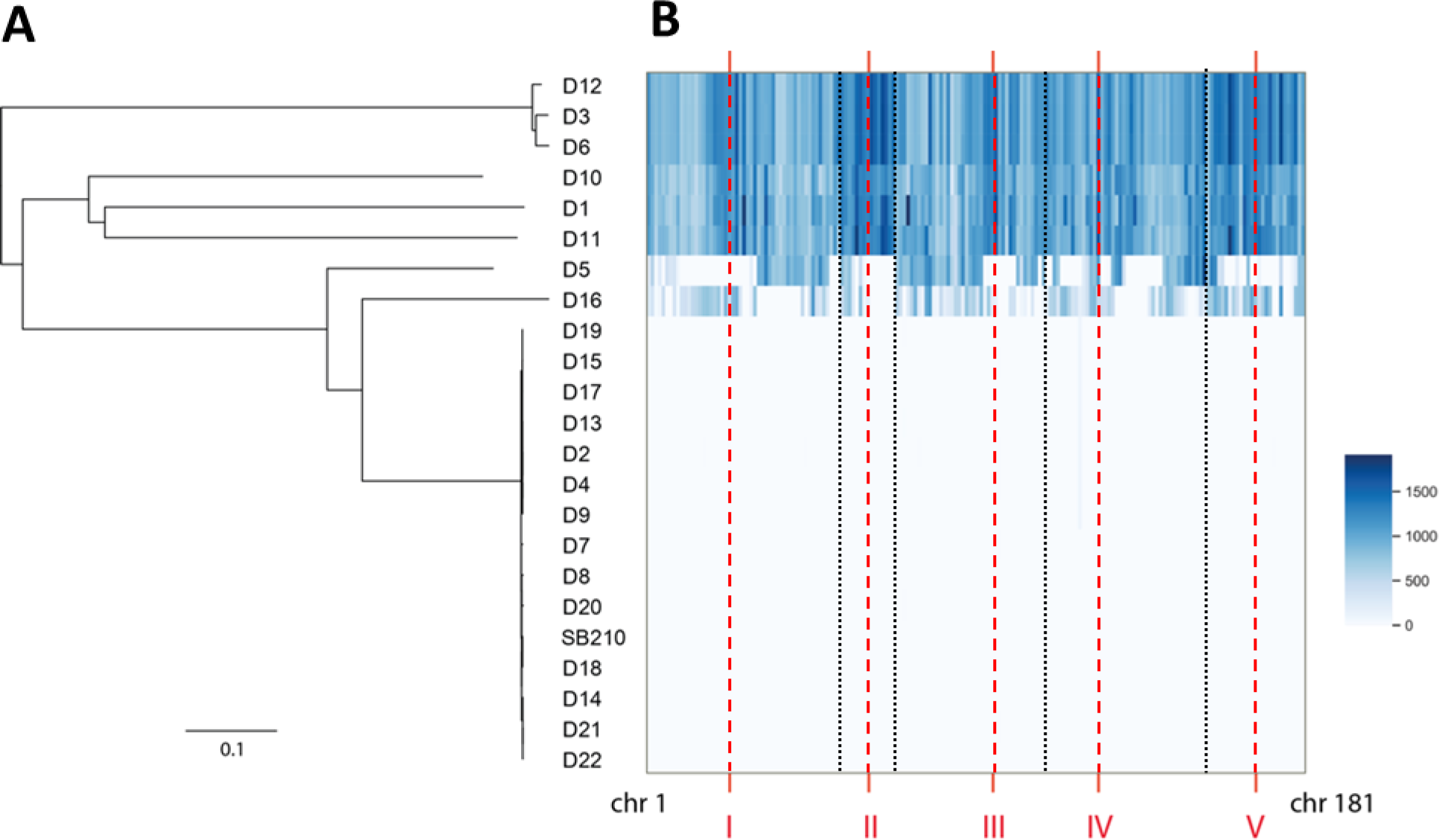
Evolutionary distances. On the left, neighbour-joining tree based on pairwise Jukes-Cantor distances between MAC genome assemblies. On the right, heatmap of the number of nucleotide differences with SB210 per 100 kb per MAC chromosome, with strains in rows and MAC chromosomes in columns (chr 1 to chr 181). The red vertical dashes indicate the approximative position of the centromeres of the five MIC chromosomes (named I to V), and the black vertical dashes the limit between these five MIC chromosomes.

Then, for each MAC chromosome individually, we reported the distances between these chromosomes and their homologs in SB210. All chromosomes of the SB210-like strains were found to be highly similar to those of SB210, while all chromosomes of D1, D10, D11 and D3/D6/D12 were distant from SB210 (Figure 2B). In contrast, the strains D5 and D16 appeared to have distinct sets of MAC chromosomes, either similar or distant from SB210. This suggests that D5 and D16 were the result of two independent and relatively recent crosses between SB210-like strains and divergent strains.

The MAC chromosomes originating from the centromeric region of the MIC chromosomes seemed to show relative higher divergences (higher number of differences with SB210 near MIC centromeres represented by dark blue hot-spots in Figure 2B, see also higher SNP density near MIC centromeres in Supplementary Material 7). This pattern could be concordant with previous observations suggesting MIC centromeres as genome innovation centres in *T. thermophila* [35].

Finally, we inferred a NJ tree from the mitochondrial genome. When mapped on the SB210 reference mitochondrial genome, the 22 Illumina datasets had an average coverage of 3,000x (Supplementary Material 8A-B), except for the extremities of this linear genome [57] that showed very high coverages, ranging from ∼25,000x and ∼40,000x (not included in analyses). SB210-like strains were grouped within the same mitotype, although D2/D4/D9/D13 have five synapomorphies. Several differences were observed between MAC and mitochondrial NJ trees: the positioning of D3/D6/D12, D5 and D16, and the grouping of D1 and D10 on the same lineage, distant from all the other strains (Supplementary Material 8C). In *T. thermophila*, the majority of mitochondria are located in the cortical ridges, with no change in their distribution during conjugation [58], as corroborated by the absence of heteroplasmy in this species [58]. Incongruences between nuclear (biparental transmission) and mitochondrial (uniparental transmission) histories are thus expected.

### Polymorphism at the MDS junctions

With the MAC genome being generated after meiosis by splicing and rearrangements from the MIC genome, we expect a background noise in SNP/indel calling at MDS junctions in a similar way recombination events do at recombination sites [22, 59]. We observed a sharp increase in the number of SNPs and indels within 20 bp of the 9,973 predicted MDS junctions: compared to the baseline polymorphism, the number of SNPs and indels increased by a factor 9 and 137 respectively (Figure 3A, values in Supplementary material 5C). This pattern was observed at both in-line and scrambled MDS junctions separately (Supplementary Material 9). Scrambled MDS refer to DNA segments that are reshuffled in the MAC in comparison with their MIC precursors [23]. Considering 20 bp around each MDS junction, the combined area of MDS junctions represented 0.4% of the MAC genome. For SB210-like strains, these areas concentrated more than half of their differences with the reference genome (49 to 68% of nucleotide differences and 70 to 74% of indels), while only a slight enrichment was observed in distant lineages (Supplementary Material 5C). These differences should almost exclusively correspond to non-homologous sequences of short IES fragments retained at MDS junctions generated by imprecise IES excision during MAC formation (Figure 1).

**Figure 3:**
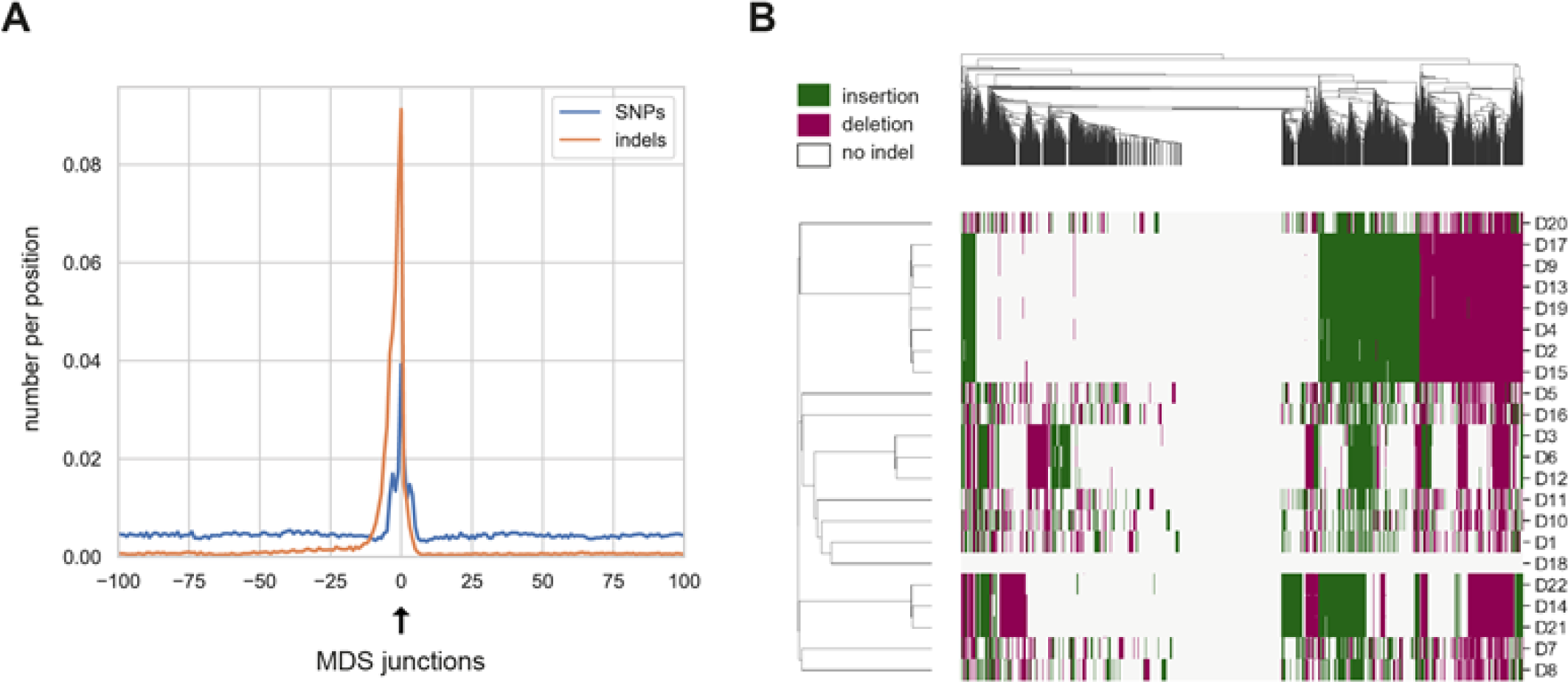
Polymorphism at the MDS junctions. **A**: numbers of SNPs and indels per position with respect to the MDS junction positions across the 22 strains. **B**: heatmap of presence/absence of indels at MDS junctions (+/− 20pb), insertions and deletions as compared with SB210 being shown in different colours. Each column represents an MDS junction. Please note that the heatmap is truncated since all the 9973 MDS cannot be represented in a small figure.

We investigated further the distribution of indels in the 22 strains. To avoid stochastic variations in their distribution across strains due to genome coverage variations, we did not use in this section the two coverage thresholds (20x and 80x) for variant calling. As shown by the heatmap in Figure 3B, most MDS junctions presented an indel in at least one strain. Across strains, we noticed three interesting patterns. Firstly, some groups of strains shared nearly identical distributions of indels at all MDS junctions: D2/D4/D9/D13/D15/D17/D19, D14/D21/D22, D3/D6/D12. Given the high rate of IES elimination errors occurring during the MAC formation, this meant that, for each of these groups, the different strains originated from the same completely assorted ancestral MAC (and hence MIC) genome. Secondly, patterns of indels at MDS junctions were found unique to these 3 groups and the remaining 9 strains (D1, D5, D7, D8, D10, D11, D16, D18, D20), even among the remaining closely related SB210-like strains (D7, D8, D20). Their patterns of indels at MDS junctions were found as distant between each other as they were from distant lineages, corroborating the fact that the variations of IES excision sites are stochastic (in theory, up to eight variants at each MDS junction, Figure 1) and not conserved between independently developing cells [59]. Thirdly, D18 had nearly identical MDS junctions to those present in the reference MAC genome of SB210, with only three indels detected at the 9,973 MDS junctions. Considering the previous observation of a very low number of nucleotide differences compared to SB210 (only 48 overall genome), we assume that the strain D18 is a recent descendent of SB210. In addition, we observed that D18 has secondarily lost its MIC genome (Supplementary material 10), as supported by microscope observations (data not shown). Amicronucleated lineages are frequent in the *Tetrahymena* genus, and can represent up to 25% of wild isolates [60].

To further illustrate the impact of imprecise IES excisions on the distribution of indels at MDS junctions in strains issued from the same conjugation event (*i.e.*, sharing the same ancestral MIC genome), we analysed the dataset published in [61]. In this study, an ancestral population was created from a single cell isolated ∼60 clonal divisions after a genomic exclusion procedure between *T. thermophila* strains SB210 and B*VII. After ∼20 clonal divisions of this ancestral population, two derived populations were created by subsampling two fractions, further grown for ∼1,000 clonal divisions. We added these three samples to our heatmap and, while they clustered together, they showed important differences in the distribution of indels (Supplementary material 11). The situation is therefore similar to our set of SB210-like strains: the three populations of [61] have the same MIC genome, but nonetheless possessed different MAC genomes due to imprecise IES excisions.

On the methodological side, we showed that SNP calling on the MAC genome of *T. thermophila* is greatly affected by the unprecise genomic rearrangements taking place during MAC formation. The fact that nearly all MDS junctions are imprecise has already been demonstrated [23, 59]. Here, we show that this general imprecision process can have profound implications for evolutionary experiments involving the MAC re-sequencing of *T. thermophila* strains (or any other ciliate species with imprecise IES excision process). If such experiment were to be conducted with a non-completely assorted ancestor or in the presence of sexual reproduction, the few dozens or hundreds of MAC-*de novo*-SNPs generated during the experiment would be diluted among more than a thousand false positive SNPs corresponding to the differences in IES excisions between the two MAC genomes (as we observed with our SB210-like strains), potentially leading to erroneous conclusions.

On a positive note, the distribution of indels at the MDS junctions creates a signature that is, given the highly imprecise nature of IES excisions and the high number of IES, unique to each MAC genome developed during the juvenile stage following sexual reproduction. This ‘barcoding’ of MAC genomes allowed us to identify in our dataset strains issued from the same developed MAC genome (*i.e.*, all chromosomes had finished their phenotypic assortment), and that our D18 strain is in fact a recent amicronucleated descendent of SB210. We therefore believe that distribution of indels at MDS junctions could become a useful tool to characterise closely related strains of *T. thermophila* with high confidence. For instance, by sequencing at a low coverage (<10x) a few hundreds of cells sampled in a given environment, it should be possible to evaluate the number of cells issued from the same MIC, but also from the same developed MAC. Those cells belonging to the same developed MAC offer high methodological facility (no need to separate MICs from MACs) and statistical power (all mutations are *de facto* MAC-*de novo*-SNPs) to estimate the time since the last sexual event, and thus the duration of asexual phase in nature. Although sexual reproduction is supposed to be frequent in natural populations of *T. thermophila* [62], its influence on population’s persistence relative to somatic selection during asexual phases is largely unknown [30]. Other eco-evolutionary perspectives would be to barcode tetrahymine cells issued from different environments using both SNPs and distribution of indels at MDS junctions to link fitness advantages (for instance better competitive or dispersal abilities) to these different sources of genomic variations.

### Recent history of strains

We further explored the very surprising observation that strains isolated by P. Doerder at different locations in the US are virtually identical, given that fifteen out of these 22 strains were previously and independently Cox1-barcoded [56, 63]. We found that only two out of the 15 common strains had the same Cox1 sequences (D10 and D11; Supplementary Material 8D). The majority of strains sharing the same MACs in our dataset did not seem representative of the natural sampling effort performed by P. Doerder. This might be due to wrong assignments and/or contaminations, at any stage of their ∼20 years of lab maintenance since their initial field sampling. Cross contaminations might explain the existence of the three groups of strains sharing the exact same distribution of indels at MDS junctions (D2/D4/D9/D13/D15/D17/D19, D14/D21/D22, D3/D6/D12). However, among the strains that were not barcoded by Doerder, which were all collected in the same geographic region (Supplementary Material 1), it is difficult to draw any conclusion on the potential cross-contaminations. For instance, the proximity between D3 and D6 could reflect a natural situation. To estimate approximately the divergence time among strains within the three groups with the same pattern of indels at MDS junctions, we looked at strain-specific mutations within each of them, which can only be MAC-*de novo*-SNPs (Supplementary Material 5D). Based on these observed numbers of strain-specific mutations, we suspect D21 and D22 on one hand, and D13 and D19 on the other hand to have split the more recently, and D3, D6 and D12 the more anciently. Comparing these numbers of MAC-*de novo*-SNPs (between 774 and 14,140 depending on the strain) with the few tens accumulated during the experiment of [61] that lasted ∼1,000 asexual divisions (∼200 days), we hypothesize that the putative cross contaminations in our dataset occurred several years ago. Unfortunately, strains were not cryopreserved nor sequenced at any stage of their lab maintenance, meaning that we cannot precisely reconstruct their recent history. Nevertheless, the overdominance of SB210-like strains suggests a massive contamination of cells issued from this reference lab strain, generally leading to replacement of the native stains. In two independent cases, an SB210-like strain probably sexually reproduced with two divergent strains (not sampled here) leading to the formation of D5 and D16.

Incongruence between our mitochondrial data and the barcoding performed in both [56] and [63] precludes any geographic assignment, and thus MAC-environment associations, from our dataset. To avoid confusion between the P. Doerder studies, *Tetrahymena* Stock Center references and our study, we hereafter only use our own denomination, D1 to D22, which matches the one we have used in all papers issued from our group since 2014 (*e.g.*, [11, 15, 39, 64, 65]).

### Single nucleotide diversity within each strain

At each position, we estimated the frequency of alternative nucleotide within each strain by normalising the number of reads supporting the difference to SB210 to a total ploidy of 47 since the sequencing was performed on DNA extractions that included both the MIC and MAC genomes (2n for the MIC genome plus ∼45n for the MAC genome). In the most distant strains, having over a million differences with SB210, most of them were homozygous (*e.g.*, 80.3% in D1; Supplementary Material 12A and values in Supplementary material 4B), while strains closer to SB210, having at most a few thousand differences, showed a much lower proportion of homozygous positions (*e.g.*, only 24.8% in strain D7; Supplementary material 12A). Indeed, the more related the strain is to the chosen reference genome, the higher the proportion of MAC-*de novo*-SNPs, and hence the proportion of unfixed positions, should compose its MAC mutational landscape (Supplementary Material 12B). A surprising pattern was to observe a higher absolute number of unfixed mutations in distant than in SB210-like strains (Supplementary Material 12C), while we expect similar numbers if mutation rates and selective pressures are the same across strains. This could reveal the presence of a methodological bias that we did not find origin and/or an impact of MIC aneuploidy (see Methods). Sequencing DNA extracts of uniquely MACs seems thus necessary to study *Tetrahymena* low frequency mutations.

### Mutational pattern

To avoid the background noise in SNP calling created by imprecise IES excisions, we estimated polymorphism metrics of the MAC genome after excluding positions within 20 bp of the MDS junctions. Since we might have missed some MDS junctions, or some MDS junctions might be placed in different positions in distant strains, we also ignored in each strain all positions within 20 bp of any indel. These criteria eliminated on average 3.7 Mb and 0.4 Mb from the genomes of distant and SB210-like strains respectively.

We first used this filtered dataset by keeping only positions for which the alternative variant as compared with SB210 had a minimum frequency of 30n (see Methods). This allowed to compare the transition/transversion ratio between a set a recently evolved strains (on which selection had less time to act) and a set of more divergent strains. Transition/transversion ratios were significantly lower in SB210-like strains compared to those observed in distant strains (0.76 and 0.96 on average for the SB210-like and distant strains respectively; Mann-Whitney U Test, p < 0.01; Supplementary Material 5E). We also observed this significant difference of ratios between distant and SB210-like strains when considering independently coding and non-coding sequences (Mann-Whitney U Test, p < 0.01; Supplementary Material 5E). This suggests the existence of a mutational bias toward transversions in *T. thermophila* with subsequent purifying selection of transversions and/or positive selection of transitions.

Secondly, we calculated the mean GC content of the MAC genomes overall strains, which was 22.3 % ± 0.01, confirming previous findings [22, 23]. When considering positions varying in only one strain (*i.e.*, the most recent mutations, that are most likely derived with respect to the ancestral state of *T. thermophila*), we observed ∼16% more G/C -> A/T than A/T -> G/C substitutions (581,592 and 502,875, respectively). This observation suggests a mutational bias toward A/T, confirming a previous, albeit more marked, report of 2.8 times more G/C -> A/T substitutions in *T. thermophila* [61, 66], in agreement with what is generally described in eukaryotes [67]. Such mutational bias potentially explains the AT-richness of *T. thermophila* genome. Altogether, our analysis of the MAC mutational pattern of *T. thermophila* strongly suggests the existence of a general bias toward G->A and C->T transitions, counterbalanced by purifying selection of these A/T mutations.

### Polymorphism metrics overall MAC genome

Across the 22 strains, the MAC nucleotide diversity measured in non-coding RNAs was found to be the lowest, followed by protein-coding sequences and finally introns and intergenic regions (Table 1). Estimations of polymorphism metrics are dependent upon the sampling, and we suspected the presence of cross-contaminations likely biasing our sampling (see section on the recent history of strains). We thus also calculated diversity metrics considering only one strain for each of the seven clusters differing in their MICs. Nucleotide diversities logically increased (‘reduced’ column in Table 1; strains D1, D3, D5, D7, D10, D11 and D16 on the basis of Figure 2). The fact that introns and intergenic regions displayed similar nucleotide diversities, whatever the dataset considered, suggests that introns are not enriched in conserved regulatory elements as it is the case in the compact genome of *Paramecium tetraurelia* [36]. Untranslated regions (UTRs), which approximately correspond in *T. thermophila* to the 200bp surrounding the genes [68], showed asymmetrical levels of polymorphism: while the 3’ UTRs displayed levels of nucleotide diversity similar to the rest of intergenic regions, we observed a marked decrease of nucleotide diversity in the 5’ UTRs (Figure 4A), suggesting that regulatory elements (*e.g.*, promoters, enhancers, silencers) are predominantly located in 5’ UTRs. Within coding regions, the nucleotide diversity was of 1.01, 0.67 and 0.23% for 4, 2, and 0-fold degenerated sites respectively for the full dataset, and 1.76, 1.17 and 0.43% for the ‘reduced’ dataset. In *Paramecium*, where species are generally distributed worldwide, these diversity values vary between 0.6-13, 0.3-8 and 0.1-1.5% respectively [36]. Given that our sample represents only a few genomic lineages that are unrepresentative of the geographic distribution of this spatially-structured species [63], we certainly underestimate the nucleotide diversity of *T. thermophila*, which could be equivalent to mid-highly diversified *Paramecium* species. It is also important to keep in mind that diversity metrics estimated with our MAC data are representative of only half of the diversity of MICs as a consequence of allelic assortment occurring during MAC development. This means that germline genomic diversity should be much higher than the one estimated here from somatic genomes.

**Figure 4:**
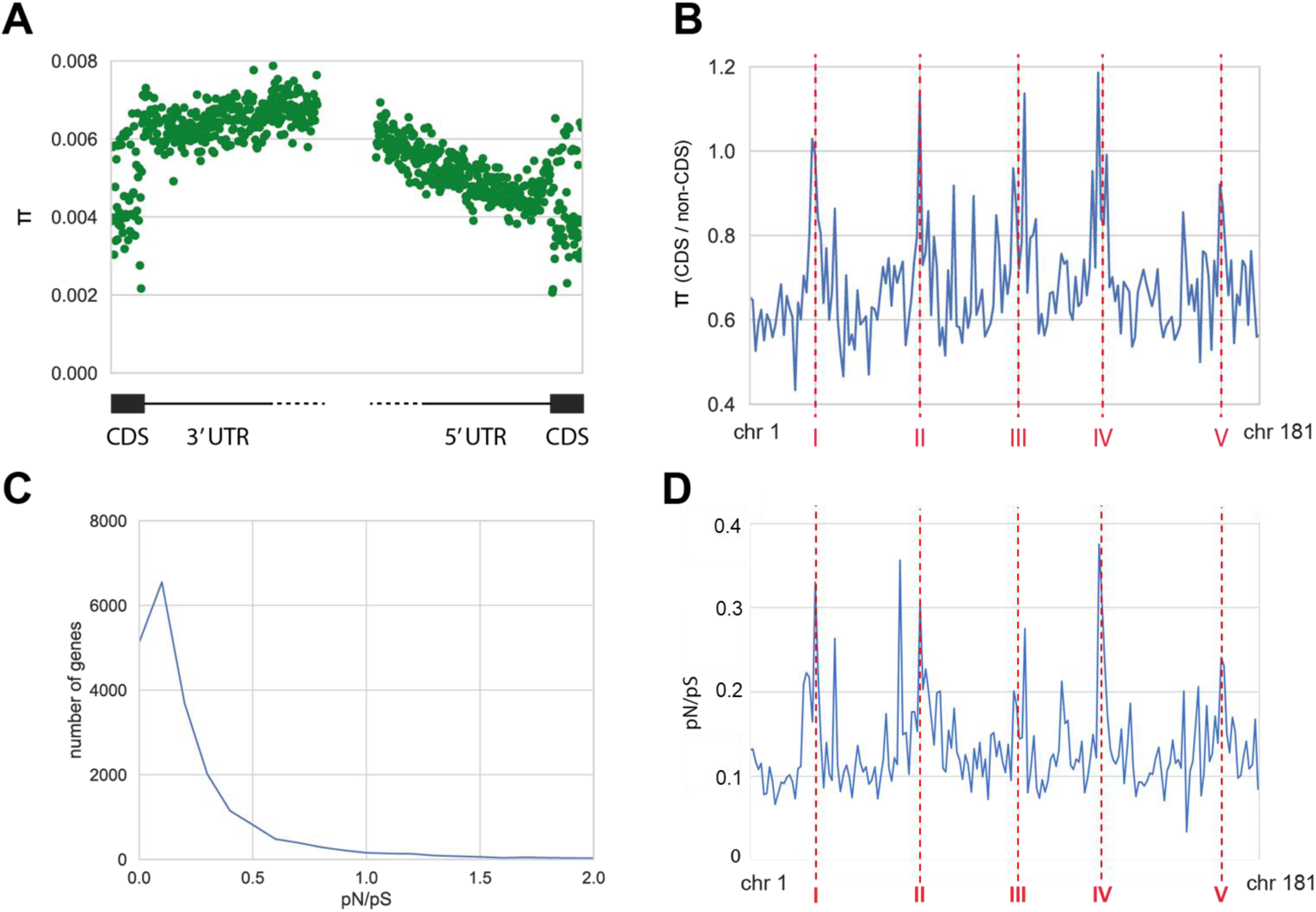
Nucleotide diversity. **A**: Nucleotide diversity (π) at the 3’ UTRs (last 50pb of the CDS and first 300pb of the intergenic region) and 5’ UTR (last 300pb of the intergenic region and first 50bp of the CDS). **B**: Ratio of π (CDS) / π (non-CDS), ‘non-CDS’ including introns and intergenic regions, for each MAC chromosome (chr 1 to chr 181 on the *x*-axis). MAC chromosomes having less than 15 genes were not included (8 chromosomes; see Supplementary material 5). The red vertical dashed lines indicate the approximative position of the centromeres of the MIC chromosome (named I to V). **C**: Distribution of pN/pS ratios. **D**: Median of genes pN/pS per MAC chromosome. The shaded area represents the confidence interval.

**Table 1:** Nucleotide diversity averages in the MAC genome. ‘reduced’ refers to a reduced taxonomic sampling of one strain per major lineage (strains D1, D3, D5, D7, D10, D11 and D16 on the basis of Figure 2), and ‘intron *’ refers to introns without their three first and three last positions.

|  | 22 strains | reduced |
| --- | --- | --- |
| ncRNA | 0.00060 | 0.00124 |
| CDS | 0.00435 | 0.00818 |
| intron | 0.00595 | 0.01170 |
| intron* | 0.00608 | 0.01202 |
| intergenic | 0.00611 | 0.01229 |
| 4-fold degenerated sites | 0.0101 | 0.0176 |
| 2-fold degenerated sites | 0.0067 | 0.0117 |
| 0-fold degenerated sites | 0.0023 | 0.0043 |

Interestingly, the difference in nucleotide diversity between the coding and non-coding sequences was heterogenous among MAC chromosomes. As shown in Figure 4B, the nucleotide diversity of coding sequences reached or slightly surpassed the nucleotide diversity of non-coding sequences in the MAC chromosomes originating from the MIC chromosome centromeres (*i.e.*, a ratio superior or equal to 1 in Figure 4B; values in Supplementary Material 5F), with a less pronounced pattern for the MIC chromosome V. Therefore, the increase of polymorphism in the MAC chromosomes originating from the MIC chromosome centromeres observed in Figure 2 appears to be principally driven by an increase of polymorphism in the coding sequences of these chromosomes.

### Polymorphism of protein-coding sequences

Considering the proportion of non-synonymous (pN) and synonymous (pS) differences in protein-coding sequences using the reduced dataset, we found that more than 85% of the protein-coding genes had a pN/pS ratio below 0.5 (Figure 4C and Supplementary Material 5G), suggesting that *T. thermophila* genes are primarily under purifying selection. We then clustered the genes by their annotation using the slim Gene Ontology (GO), and reported in Supplementary Material 13 the distribution of pN/pS values for each slim GO term. Focusing on the ‘cellular component’ GOs, the 78 genes encoding for ‘proteins of the cilium’ had the lowest median pN/pS value (0.02). We hypothesize the constraints on these genes are due to the fundamental role of cilia in feeding and locomotion processes in ciliates, and the general stability of cilia organization within the ‘morphospecies’ of the genus *Tetrahymena* [69, 70]. The 62 genes encoding for ‘extracellular proteins’ had the highest median pN/pS value (0.19) followed by the 17 genes encoding for ‘extracellular space’ (median pN/pS = 0.17), and the 43 genes encoding for ‘plasma membrane proteins’ (median pN/pS = 0.12). This suggests that many proteins interacting with the extracellular environment (membrane receptors, effectors, etc.) accumulate mutations faster than the rest of the genome. *T. thermophila* is typically found in floating or emergent vegetation near the shore of small ponds, including roadside ditches, and sometimes downstream from ponds [70]. These small habitats can have, for instance, very diverse and fluctuating chemical composition, thermal regime or biotic composition. Rapid adaptation of proteins interacting with such a variety of extracellular environments are probably key to ensure long-term persistence of the species. Across all categories, the only GO term with a notably high median pN/pS value was ‘cell adhesion’ (category ‘biological process’; 86 genes; median pN/pS = 0.24), which was dominated by genes encoding metalloproteases (Leishmanolysin family proteins). Some strains of *T. thermophila* present cooperative behaviours: cells aggregate to exchange growth factors when environment degrades [71], kin recognition presumably modulating aggregative behaviour to exclude cheaters issued from non-cooperative strains from social interactions [16]. We speculate that competition between cooperative and non-cooperative strains contributes to the rapid evolution of cell adhesion proteins in *T. thermophila*.

We then compared the polymorphism of *T. thermophila* to the divergence with other *Tetrahymena* species using the Direction of Selection (DoS) metric. We found that 82.4% of genes had negative DoS values (mean=-0.0897, Supplementary Material 5H), suggesting that slightly deleterious mutations are generally segregating. As for pN/pS values, we calculated the distribution of DoS values for each slim GO term (Supplementary Material 14). Intriguingly, the highest values of the distribution were found for ‘ribosomes’ (median value of 0.023) and ‘structural constituent of ribosomes’ (median value of 0.034), suggesting positive selection on genes involved in ribosomal function distinguishing *T. thermophila* from its relatives of the genus *Tetrahymena*. On the opposite, the lowest DoS values were found for ‘extracellular space’ (median value of −0.134), despite especially high pN/pS value (see above). This means that an important fraction of the numerous non-synonymous SNPs found in these genes might be slightly deleterious and still segregating in *T. thermophila*, and/or that selection is relaxed in *T. thermophila* compared with its relatives.

We calculated for each MAC chromosome the median pN/pS and DoS values of all genes (Supplementary Material 5I and 5J). As shown in Figure 4D, the MAC chromosomes originating from the centromeres of the MIC chromosomes presented the highest median pN/pS. This result indicates that the relative increase of nucleotide diversity mentioned above in these CDS (Figure 4B) was due to an increase in non-synonymous mutations. As for nucleotide diversity (Figure 4B), the MAC chromosome originating from the centromere of the MIC chromosome V displayed a moderate increase compared to the other four centromeric regions. We also showed a sharp decrease in the number of genes for which the DoS could be calculated in MIC centromeric regions (Supplementary Material 15). Only 49% of genes present in the MAC chromosomes originating from the MIC centromeres had DoS values while 67% of genes present in the rest of the genome had DoS value. Missing DoS genes corresponded to genes for which orthologs were not found in the other species of the *Tetrahymena* genus, *i.e.*, recent (duplicated) and/or rapidly evolving genes. For genes with identifiable orthologs in MIC centromeric regions, DoS were notably low (−0.1191) compared with the rest of the genome (−0.0897, Supplementary Material 15), suggesting distinct evolution for the conserved genes present in these regions.

Altogether, results suggest a two-speed evolution of MIC pericentromeric regions. Firstly, these regions act as diversification centres hosting more recent and/or fast evolving genes than the rest of the genome with high pN/pS values, confirming previous results [35]. Contrary to many taxa (animals, plants) presenting both symmetric (male) and asymmetric (female) meiosis, ciliates only undergo female meiosis: in both parental cells, three of the four meiotic products are eliminated. Such competition between meiotic products confers a selfish advantage to chromosomes’ centromeres that increase their probability of transmission through, *e.g.*, DNA sequences helping in the recruitment of centromeric proteins. This competition leads to the so-called “centromere-drive” [72]. Fast evolution of *T. thermophila* MIC centromeres could in part result from this inter-chromosome competition, which does not seem mitigated by well-known suppressors (*e.g.*, [73]). Secondly, genes from MIC centromeric regions that are conserved across the *Tetrahymena* genus evolve especially slowly compared with the rest of the genome. This could be explained by the presence of key structural genes preserving cell integrity and/or centromere structures in these regions.

We finally tested if the genes around MIC centromeric regions presenting higher pN/pS, dN/dS and DoS values than the means calculated on these regions had specific functions. After controlling that results were not driven by a genome-wide pattern, we found that DoS provided no obvious tendency. However, pN/pS (intraspecific divergence) revealed a slight enrichment tendency around MIC centromeres toward genes ensuring protein binding, carboxypeptidase activity, exopeptidase activity, phosphorylation processes, and kinase activity. For dN/dS (interspecific divergence), the same slight tendency for phosphorylation processes and kinase activity was observed. This means that genes present in MIC centromeric regions involved in protein binding, carboxypeptidase and exopeptidase activity are evolving rapidly only in *T. thermophila*. Carboxypeptidase and exopeptidase activity, which are involved in protein degradation, notably in the digestion process. Further investigation will be required to determine if such increased diversity relates to the artificial diet imposed to *T. thermophila* since years after its isolation from natural ponds. On the contrary, genes involved in phosphorylation and kinase activity are evolving similarly faster in *T. thermophila* as for the rest of the tested members of the genus *Tetrahymena*. Interestingly, genes with no DoS around MIC centromeres (*i.e.*, no orthologs in the *Tetrahymena* genus) were enriched toward protein binding, kinase activity and phosphorylation processes. This suggests both recent bursts and rapid evolution of these genes usually involved in the transduction of signals and in protein and/or transcription regulation (*e.g.*, [74]). Such a fast evolution is thus expected. In the future, it would be interesting to determine if these genes display the same MIC pericentromeric location and high burst rate in other *Tetrahymena* species.

## Conclusion

We provide here the first intraspecific genomic dataset in the model ciliate *T. thermophila*. While we could describe several interesting patterns, this work should be seen as a first step. Increasing the number of complete *T. thermophila* MAC, but also MIC, genomes with long read methods should help to obtain more precise assemblages. This would allow to tackle questions that are still open in both mechanistic biology and ecology. Especially, one strong limitation to our study is the impossibility to assign strains to their geographic origin, which prevented us to explore genome/environment associations in this model species. Another limitation is our small number of diverging strains (seven). While we could provide a first assessment of MAC intraspecific diversity in *T. thermophila*, we clearly underestimate its true diversity. One the one hand, our ‘reduced’ sample is probably not a good representative of the species, and on the other hand, MACs host only half of MIC diversity (as half of alleles are lost through phenotypic assortment following MAC formation). This means that basic population genetic parameters such as heterozygosity or Hardy-Weinberg equilibrium of MICs cannot be estimated by the sequencing of MACs. However, the unfortunate discovery of half of the studied strains sharing the same MIC as SB210, the other being distant, offers an unexpected opportunity to explore phenome/genome associations. Using our 22 MAC landscapes, it is possible to explore the links between (epi)genetic and phenotypic diversity in a sample for which many ecologically relevant traits have been measured (*e.g.*, [11, 39, 40]).

## Supporting information

Supplementary Material 1 Derelle et al.

Supplementary Material 2 Derelle et al.

Supplementary Material 3 Derelle et al.

Supplementary Material 4 Derelle et al.

Supplementary Material 5 Derelle et al.

Supplementary Material 6 Derelle et al.

Supplementary Material 7 Derelle et al.

Supplementary Material 8 Derelle et al.

Supplementary Material 9 Derelle et al.

Supplementary Material 10 Derelle et al.

Supplementary Material 11 Derelle et al.

Supplementary Material 12 Derelle et al.

Supplementary Material 13 Derelle et al.

Supplementary Material 14 Derelle et al.

Supplementary Material 15 Derelle et al.

## Acknowledgements

We thank Nicolas Schtickzelle for having provided the strains used in this study and for fruitful interactions about unpublished genetic datasets he obtained in his lab. We also thank Léonard Dupont for his help in Python programming.

## Author contribution

R.D.: conceptualization, methodology, selection and formal analysis, writing - original draft, writing - review and editing

R.V.: methodology, writing - review and editing

S. J.: writing - review and editing

M. H.: strain maintenance, molecular biology work

I. A.: bioinformatic resource, writing - review and editing

H. P.: conceptualization, funding acquisition, bioinformatic resource, methodology, formal analysis, writing-review and editing

D. L.: conceptualization, funding acquisition, selection analyses, molecular biology work, writing - original draft, writing - review and editing

