## Supplementary Material 2 Derelle et al. for "The macronuclear genomic landscape within *Tetrahymena thermophila*"

**Supplementary Material 2: DNA extraction electrophoresis**

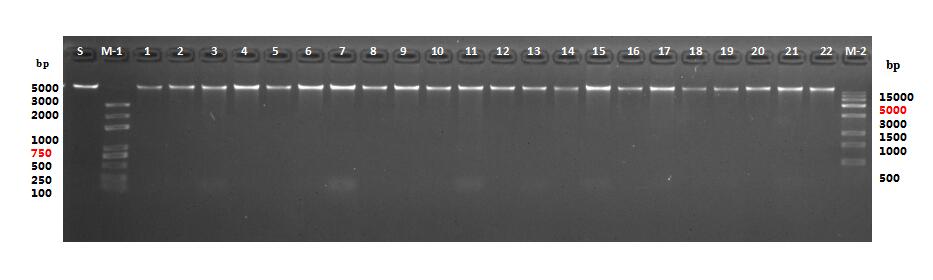

Electrophoresis of the 22 DNA extractions before sequencing. S is a positive control of DNA extraction, M-1 and M-2 are ladders, and strains D1 to D22 are indicated by the 1 to 22 numbers.
