## Supplementary Material 3 Derelle et al. for "The macronuclear genomic landscape within *Tetrahymena thermophila*"

**Supplementary Material 3: bioinformatics procedures**

**Modifications of the reference MAC SB210 genome assembly**

The genome assembly of *T. thermophila* strain SB210 published in (Sheng, et al. 2020) was downloaded from <http://ciliate.org>. The assembly was first polished using Pilon v1.23 (Walker, et al. 2014) and the Illumina reads published in the same study (SRA database: SRR11906179) to correct potential sequencing/assembly errors. The two rounds of polishing generated 4027 and 55 modifications of the assembly (mostly one nucleotide indels).

We then renamed the 181 MAC chromosomes according to their respective origin in the 5 MIC chromosomes. The MAC chromosomes were mapped onto the MIC genome assembly [ref] downloaded from <http://ciliate.org> using minimap2 v2.17 (Li 2018) with the parameter ‘-x asm5’. For each MAC chromosome, the position in the MIC genome was defined as the centre of their mapping hits. The MAC chromosomes were finally named from ‘chr_001’ to ‘chr_181’ given these positions (chr_001 corresponding to the MAC chromosome located at the 5’ end of the MIC chromosome 1). Finally, MIC chromosomes being centromeric, the MAC chromosomes corresponding to the MIC centromeres were identified by summing up the length of all MAC chromosomes of a given MIC chromosome as in (Hamilton, et al. 2016).

This corrected MAC genome assembly is referred in the text to as ‘the SB210 genome assembly’.

**Creation of a new predicted gene set**

We noticed during our preliminary analyses that many of the genes defined in (Sheng, et al. 2020) were suspicious (e.g. presence of stop codons after genome polishing, unexpected large number of long introns). Therefore, we decided to create a new gene set, based mostly on the high quality Ensembl gene prediction (v2.2.48; 24,725 protein-coding genes and 927 ncRNA genes).

Firstly, the sequences of Ensembl protein-coding genes that did not have any ambiguous nucleotide (i.e. no ‘N’ in their sequence; 24,152 genes) were Blasted on to the SB210 genome assembly with a 98% identity threshold (Blastn; v2.2.28). For each gene, the Ensembl prediction was transferred to the SB210 genome assembly using a custom python script if (i) the Blast hit covered the entire gene, (ii) there was no indel between the Blast hit and the gene sequence, (iii) the inferred exons did not overlap with previously defined exon positions and (iv) the predicted protein sequence inferred from the inferred exons did not contain any internal stop codon. These criteria led to the transfer of 23,643 protein-coding gene annotations to the SB210 genome assembly.

We then analysed the annotations of the protein-coding genes defined in (Sheng, et al. 2020) using two distinct approaches. Firstly, we transferred to the SB210 genome assembly the annotation of the genes with a name starting with ‘TTHERM_’ (e.g. genes initially defined by Ensembl) if (i) the inferred exons did not overlap with previously defined exon positions and (ii) the predicted protein sequence inferred from the inferred exons did not contain any internal stop codon. This step led to the transfer of 2,270 protein-coding gene annotations. Secondly, we transferred to the SB210 genome assembly the annotation of the genes with a name starting with ‘g’ (e.g. genes newly defined in (Sheng, et al. 2020); 919 genes) if they satisfied the two previous criteria and if their protein sequence had a blast hit (Blastp; v2.2.28) with a e-value below 1e-5 against the predicted proteome of either *Paramecium tetraurelia*, *Oxytricha trifallax* str. JRB310 or *Ichthyophthirius multifiliis* (alldownloaded from Ensembl Protist; <https://protists.ensembl.org>). This step led to the transfer of 105 protein-coding gene annotations, and a final gene set of 26,018 protein-coding genes.

Finally, the sequences of Ensembl ncRNA genes were Blasted on to the SB210 genome assembly with a 98% identity threshold (Blastn; v2.2.28). For each gene, the Ensembl prediction was transferred to the SB210 genome assembly if the Blast hit covered the entire ncRNA gene and did not overlap with any coding exon, leading to the annotation of 546 ncRNA genes.

**Identification of MDS junctions**

The corrected SB210 MAC genome assembly was aligned to the SB210 MIC genome assembly (Hamilton, et al. 2016), downloaded from <http://ciliate.org>, using minimap2 v2.17 (Li 2018) with the parameter ‘-x asm5’. Then, an in-house python script identified MDS junctions from the minimap2 output as follows:

_ only alignments with a score equal to 60 (i.e. maximal values) were considered for further analyses.

_ in-line MDS junctions were identified within alignments, corresponding to insertions in the MIC genome of at least 200 non-N nucleotides.

_ scrambled MDS junctions were identified between alignments, when (i) the positions of the extremity of two MAC alignments were separated by less than 10 bp (the position of the MDS junction corresponding to the average of the two extremity positions) and (ii) the corresponding MIC segment between the two extremities contained at least 200 non-N nucleotides.

A total of 9,973 MDS junctions were identified by this approach, including 7,273 ‘in-line’ MDS junctions (*i.e.*, the two MDS were adjacent and on the same strand in the MIC genome) and 2,700 ‘scrambled’ MDS junctions (*i.e.*, the two MDS were joined following a genomic rearrangement). This number of scrambled MDS junctions was very similar to that observed in (Sheng, et al. 2020) (2,711 scrambled MDS junctions). Over the 7,544 MDS junctions identified by (Hamilton, et al. 2016), 6,911 were found within 10-bp of one of the MDS junctions identified in this study. We thus consider our dataset reliable to assess intraspecific polymorphism patterns at MDS junctions.

**Command lines for read mapping and SNP/indel calling**

The program versions are indicated in Material and methods. The following command lines were repeated for each strain (D1 to D22). Firstly, trimmed reads were aligned to the reference genomes (MAC and mitochondrial genomes) and the alignments were filtered to retain high quality matches (i.e. with a mapping quality of at least 30):

- bwa mem -t 4 reference_genome.fa D1_trimmed.fq.gz D1_2_trimmed.fq.gz > D1.sam
- samtools view -q 30 -h -b -S D1.sam > D1_filtered.bam
- samtools sort -m 19G -o D1_sorted.bam D1_filtered.bam

Then to obtain coverages:

- bedtools genomecov -ibam D1_sorted.bam -d > D1_coverages.txt

Then to obtain SNPs and indels:

- samtools mpileup -t DP -uvf reference_genome.fa D1_sorted.bam > D1.vcf
- bcftools call -f GQ -o D1_snps.vcf -O v -V indels -cv D1.vcf
- bcftools call -f GQ -o D1_indels.vcf -O v -V snps -cv D1.vcf
- vcfutils.pl varFilter -in D1_snps.vcf > D1_snps.filt.vcf
- vcfutils.pl varFilter -in D1_indels.vcf > D1_indels.filt.vcf
