## Supplementary figures and images for "The macronuclear genomic landscape within *Tetrahymena thermophila*"

### Supplementary Material 4 Derelle et al.

**Supplementary Material 4: Gene counts per MAC chromosome**

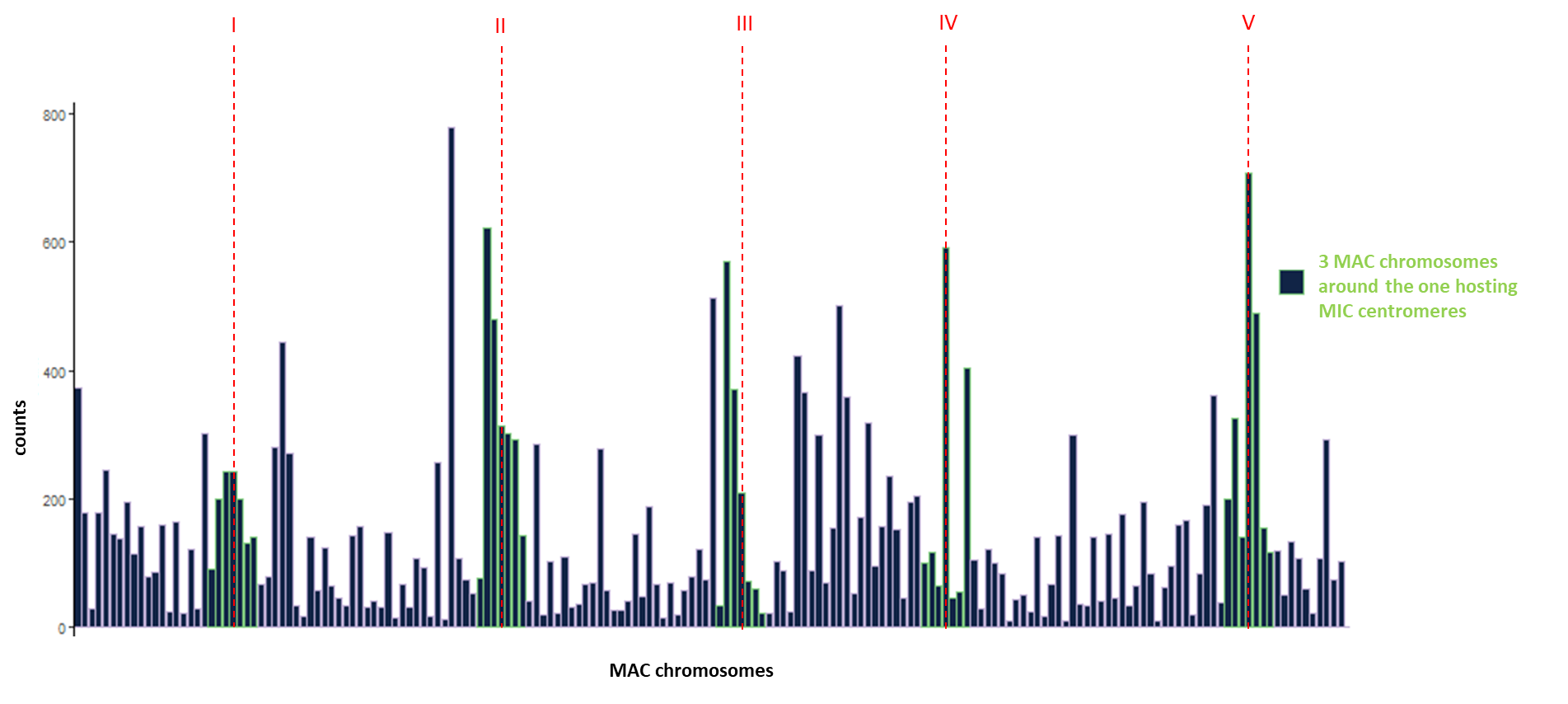
