## Supplementary Material 6 Derelle et al. for "The macronuclear genomic landscape within *Tetrahymena thermophila*"

**Supplementary Material 6: Neighbour-Joining tree**

This analysis was based on the 22 MAC genome assemblies generated in this study (described in Supplementary Material 3) and MAC genomes assemblies downloaded from <http://ciliate.ihb.ac.cn>.

The genomes were analysed using andi v0.13 using default parameters (Haubold et al., 2015), and a neighbour-joining tree was built from the resulting kmer-based distances using NINJA v1.2.2 (Wheeler, 2009).

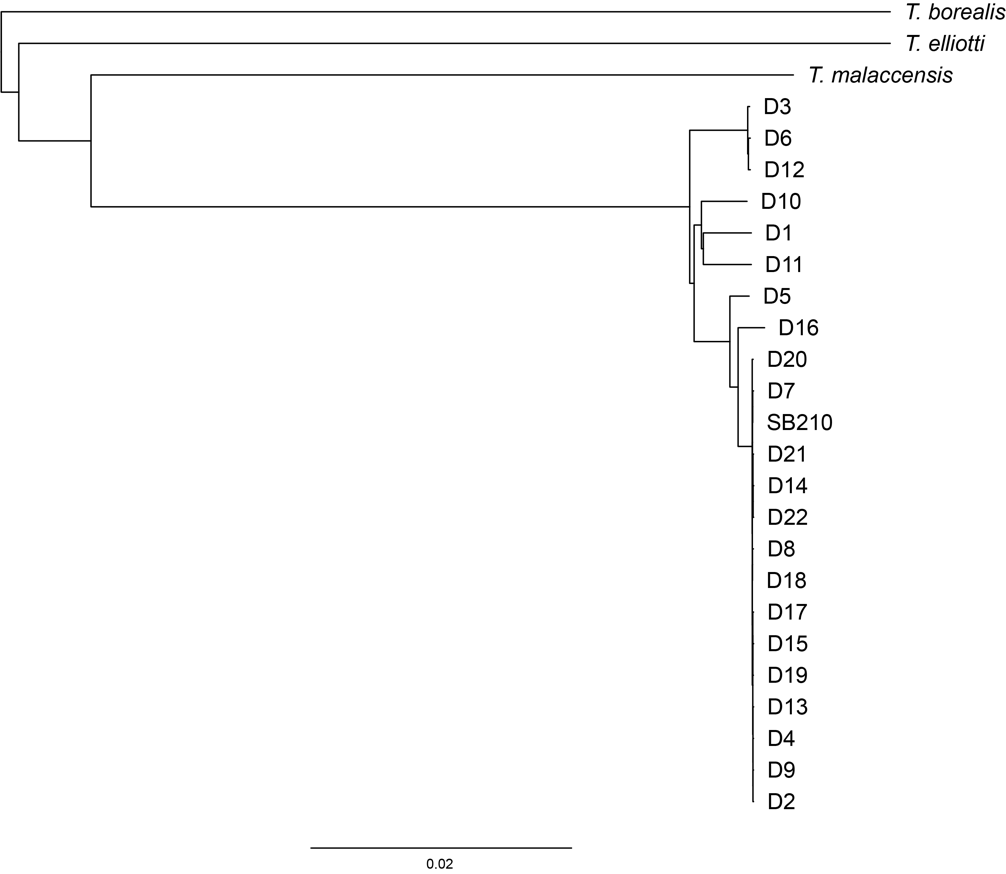

Haubold, B., Klotzl, F., and Pfaffelhuber, P. (2015). andi: fast and accurate estimation of evolutionary distances between closely related genomes. Bioinformatics *31*, 1169-1175.

Wheeler, T.J. (2009). Large-scale neighbor-joining with NINJA. In S.L. Salzberg and T. Warnow (Eds.), Proceedings of the 9th Workshop on Algorithms in Bioinformatics. WABI 2009, 375-389.
