## Supplementary Material 7 Derelle et al. for "The macronuclear genomic landscape within *Tetrahymena thermophila*"

**Supplementary Material 7: Number of SNPs per 100,000 nucleotides per MAC chromosome (overall strains)**

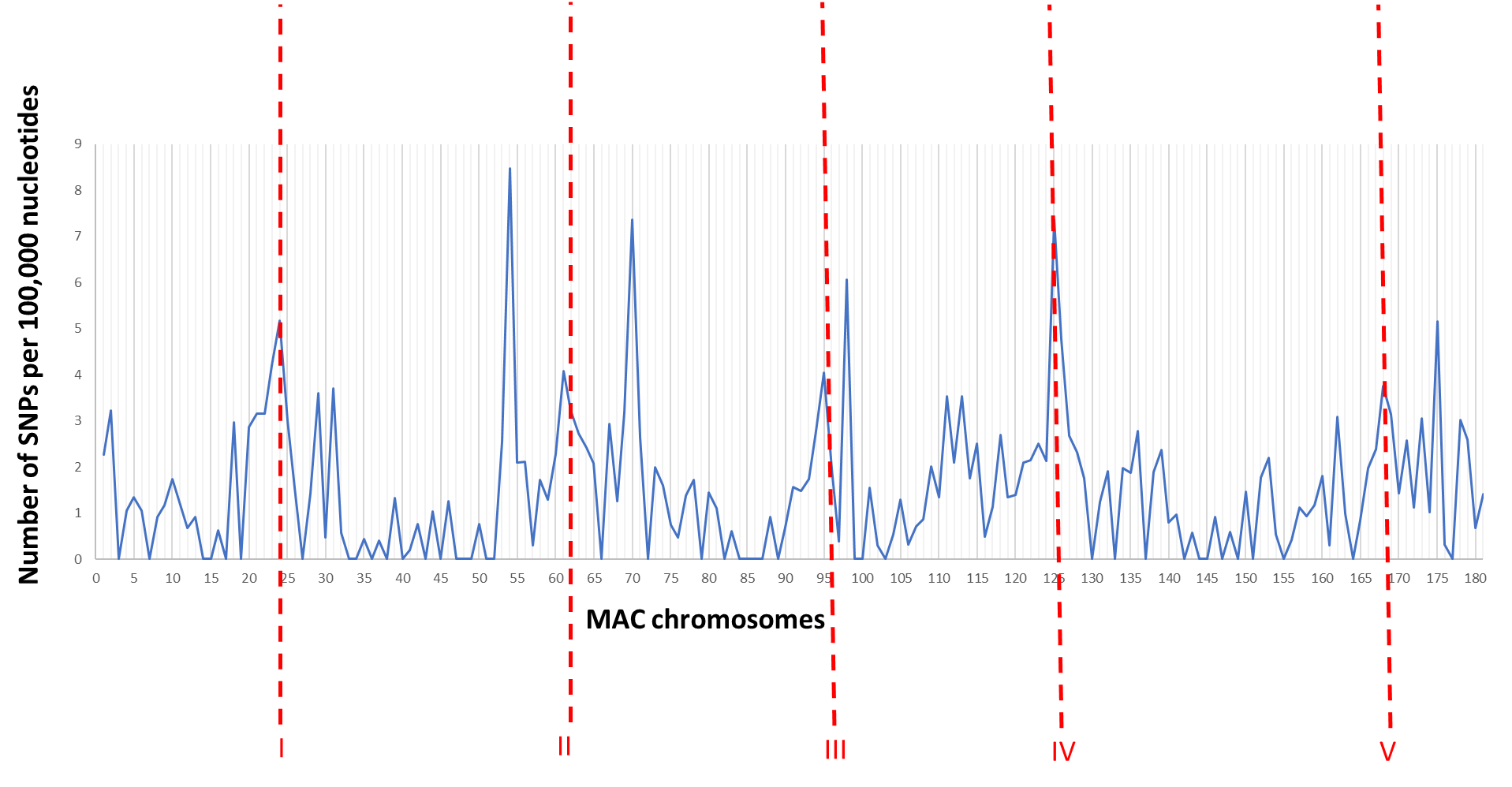

The positions of MIC centromeres are represented by the five red dashed lines.
