## Supplementary Material 8 Derelle et al. for "The macronuclear genomic landscape within *Tetrahymena thermophila*"

**Suplementary material 8: mitochondrial genomes**

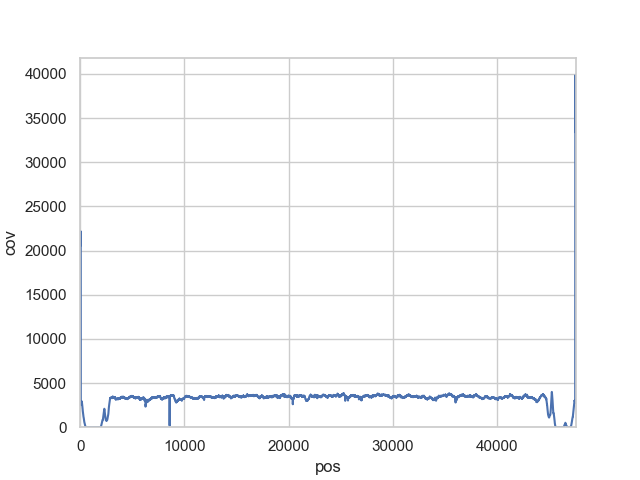

A)

Average coverage along the mitochondrial genome over the 22 *T. thermophila* strains. The 2kb from each extremity, composed of repeat regions, were excluded.

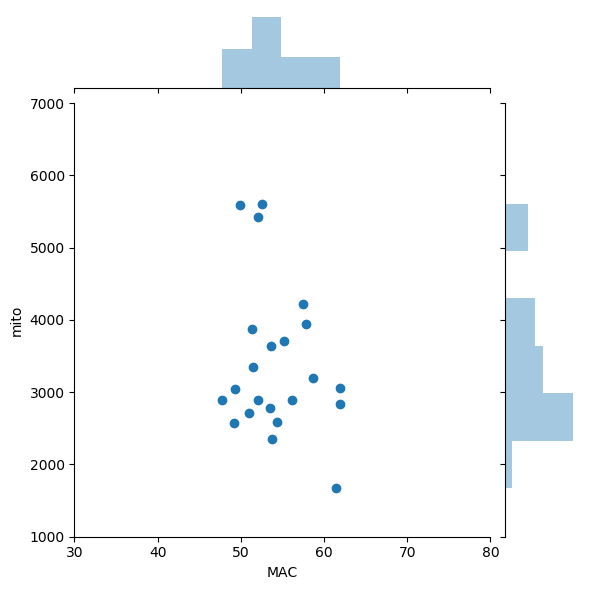
B)

za

Mitochondrial coverage as a function of MAC coverage in the 22 strains, with respective distributions on the top and the right of the graph.

C)

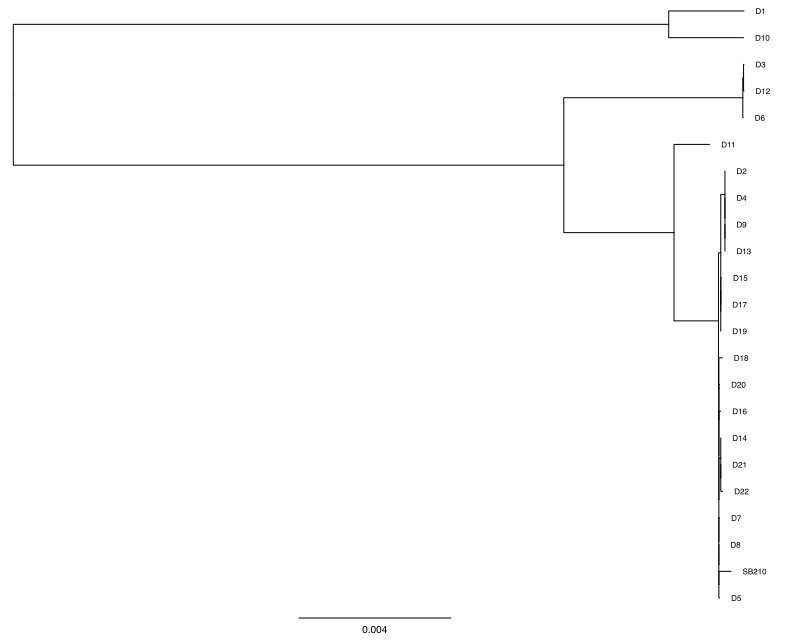

Neighbour-joining tree based on the whole mitochondrial genome of our 22 strains plus the reference SB210 mitochondrial genome.

D)

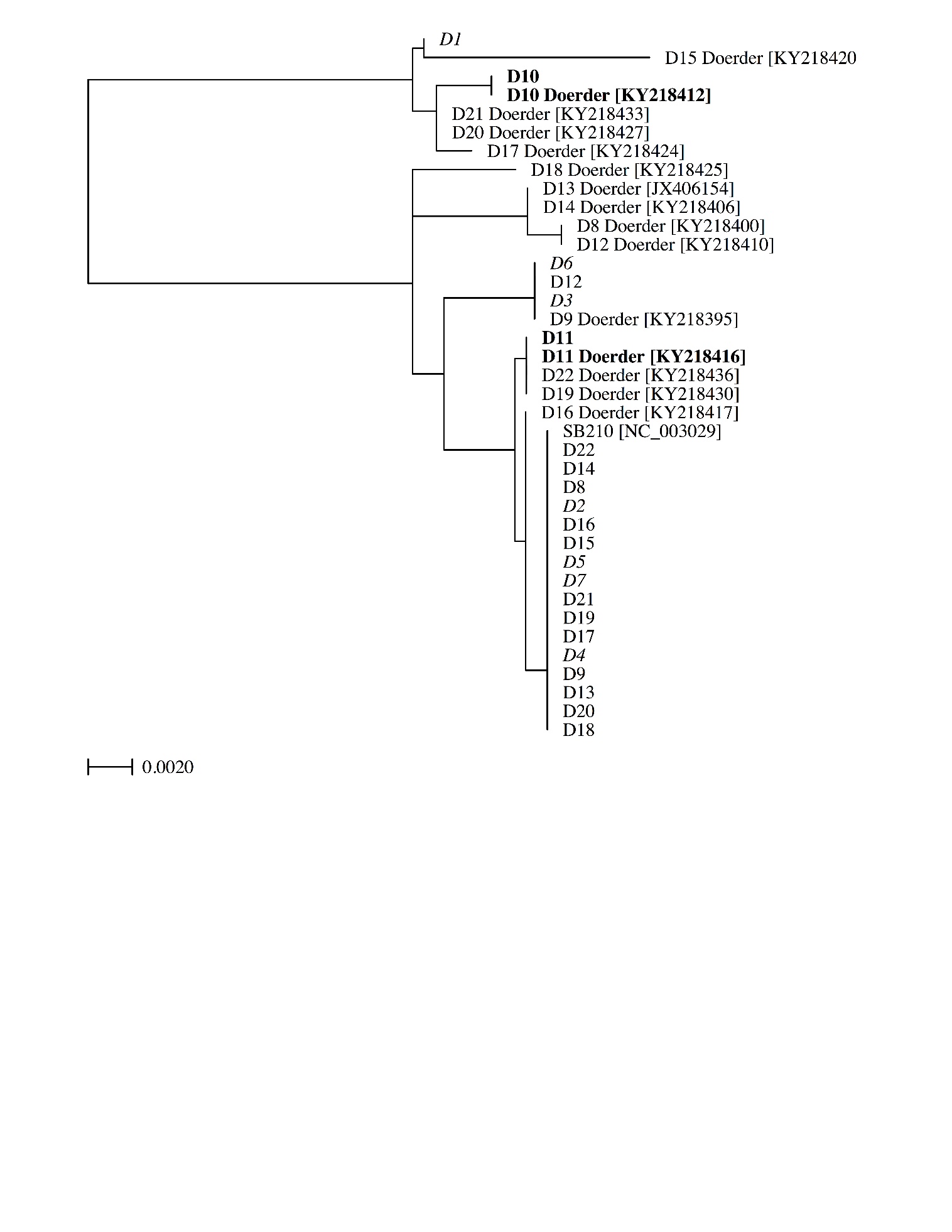

Neighbour-joining tree based on based on the Cox 1 region uniquely of our 22 strains plus the reference SB210 mitochondrial genome and the 15 common sequences between our work and those of Zufall et al. (2014) and Doerder (2019). Strains in bold indicate congruence between the two studies, strains in italic were not sequenced before our study.
