## Supplementary Material 9 Derelle et al. for "The macronuclear genomic landscape within *Tetrahymena thermophila*"

**Supplementary Material 9: polymorphism around MDS junction**

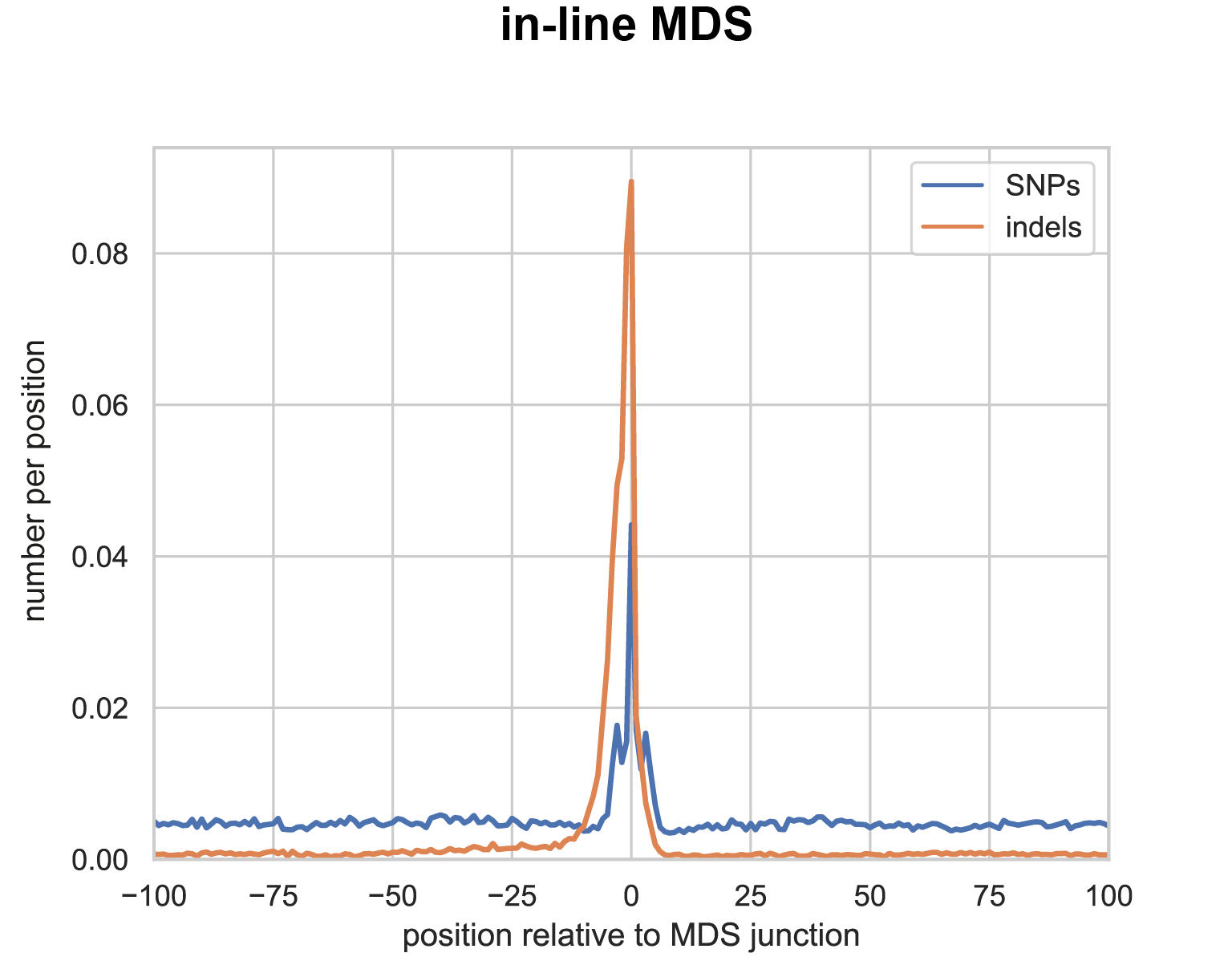

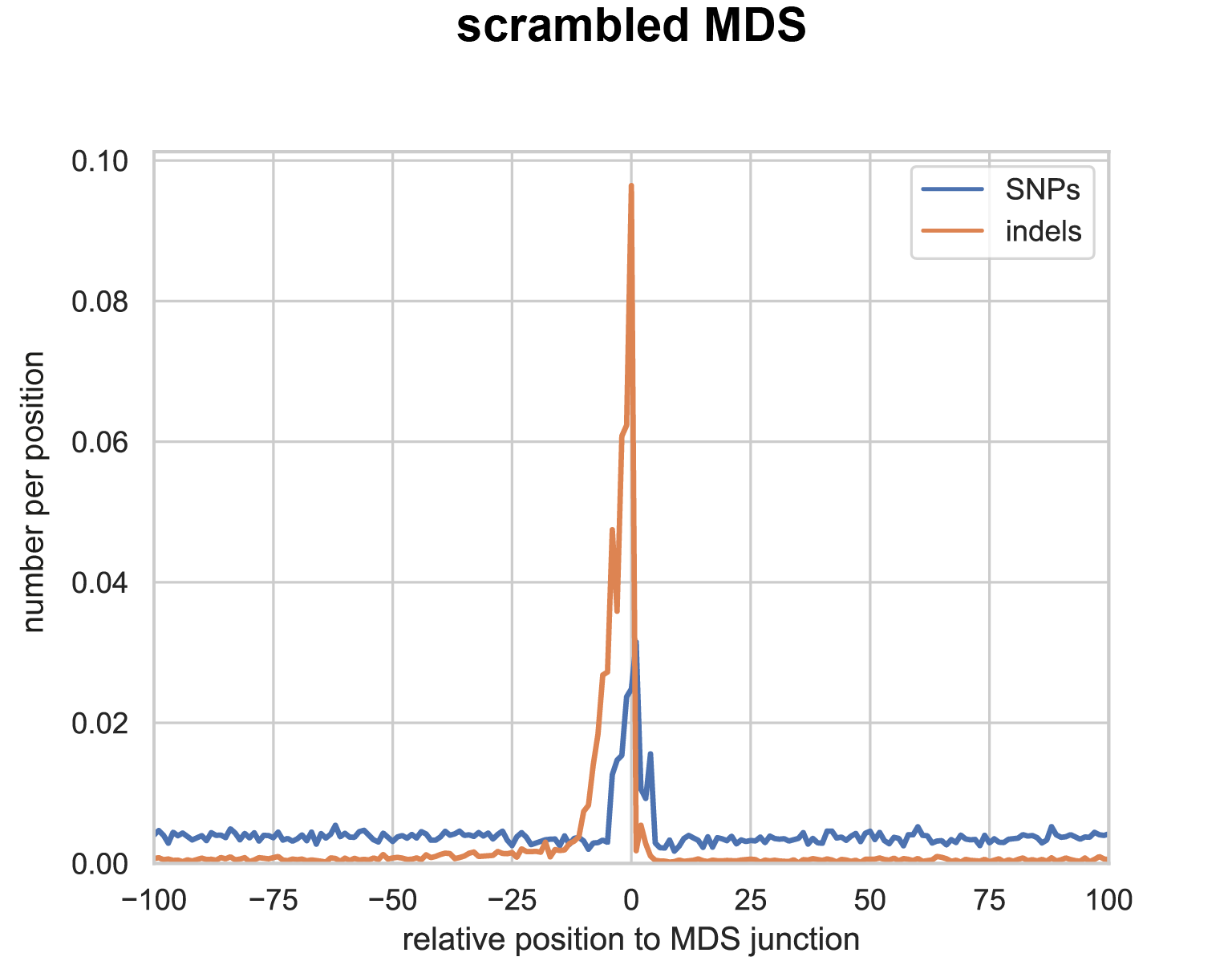

Number of SNPs and indels per position with respect to the MDS junction positions for in-line and scrambled MDS across the 22 strains
