## Supplementary Material 10 Derelle et al. for "The macronuclear genomic landscape within *Tetrahymena thermophila*"

**Supplementary Material 10: D18**

This document attempts to demonstrate, using genomic evidences, that the strain D18 is amicronucleated (*i.e.*, absence of MIC genome). The strategy here consisted on blasting the IES (*i.e.*, sequences usually specific to the MIC genome) on the genome assembly of the different strains, which should contain both the MIC and MAC genomes, to identify the presence or absence of MIC genome.

Firstly, the trimmed reads of each of the 22 strains were assembled into contigs with SPAdes v3.14.1 (Prjibelski et al., 2020) using the options ‘-k 105 --only-assembler –careful’. The assembly statistics are provided in the following table:

| strain | size (Mb) | N50 (kb) | nb seq | longest seq (kb) | % GC | % N |
| --- | --- | --- | --- | --- | --- | --- |
| D1 | 108.7 | 325.82 | 1276 | 1534.3 | 22.3 | 0.005 |
| D2 | 109.2 | 275.51 | 3231 | 1545.53 | 22.3 | 0.0077 |
| D3 | 113 | 356.79 | 1380 | 2096.37 | 22.32 | 0.0042 |
| D4 | 108.8 | 270.83 | 3249 | 1294.23 | 22.3 | 0.0057 |
| D5 | 114.4 | 343.67 | 1480 | 1635.37 | 22.32 | 0.0056 |
| D6 | 109.7 | 370.36 | 1237 | 1935.81 | 22.31 | 0.0047 |
| D7 | 115.4 | 317.15 | 1541 | 1336.47 | 22.32 | 0.0056 |
| D8 | 113.2 | 270.11 | 3718 | 1295.31 | 22.31 | 0.0053 |
| D9 | 108.8 | 280.64 | 3207 | 1294.2 | 22.3 | 0.0067 |
| D10 | 111.4 | 279.25 | 1423 | 1922.01 | 22.31 | 0.0047 |
| D11 | 111.7 | 271.28 | 1360 | 1639.3 | 22.31 | 0.0051 |
| D12 | 110.2 | 334.82 | 1304 | 1638.83 | 22.31 | 0.0054 |
| D13 | 107.8 | 316.38 | 1742 | 1294.21 | 22.31 | 0.0064 |
| D14 | 108 | 301.85 | 2238 | 1415.04 | 22.3 | 0.0066 |
| D15 | 110.6 | 269.71 | 3670 | 1056.55 | 22.32 | 0.005 |
| D16 | 108 | 328.09 | 2204 | 1645.06 | 22.3 | 0.0055 |
| D17 | 110.1 | 286.75 | 3255 | 1294.17 | 22.33 | 0.0071 |
| D18 | 103.7 | 345.13 | 998 | 1589.16 | 22.29 | 0.0049 |
| D19 | 108.6 | 319.58 | 2045 | 1296.67 | 22.3 | 0.0057 |
| D20 | 107 | 313.06 | 1268 | 1293.21 | 22.29 | 0.0053 |
| D21 | 107 | 332.93 | 1497 | 1426.63 | 22.3 | 0.0064 |
| D22 | 107.3 | 305.31 | 2044 | 1501.08 | 22.29 | 0.0072 |

We then extracted the 7,544 IES (28,529,842 positions) from the SB210 MIC genome (Hamilton et al., 2016) using an in-house script, and blasted this set of sequences on each of the 22 genome assemblies (blastn v2.6.0; options ‘-perc_identity 98 -evalue 1e-100’).

Finally, we reported in the following table, for each strain, the number of IES having a blast hit against the genome assembly and the number of distinct IES positions matching the genome assembly:

| strain | nb IES | nb IES positions |
| --- | --- | --- |
| D1 | 1828 | 1123173 |
| D2 | 1350 | 2666430 |
| D3 | 2645 | 1770039 |
| D4 | 1299 | 2535631 |
| D5 | 4152 | 4062190 |
| D6 | 1828 | 1268319 |
| D7 | 4894 | 5306020 |
| D8 | 2157 | 4299870 |
| D9 | 1330 | 2579992 |
| D10 | 2614 | 1646480 |
| D11 | 2751 | 1809949 |
| D12 | 2286 | 1430468 |
| D13 | 1299 | 2068254 |
| D14 | 1380 | 2327504 |
| D15 | 1522 | 3019333 |
| D16 | 1042 | 1251402 |
| D17 | 1503 | 2900962 |
| **D18** | **122** | **35134** |
| D19 | 1512 | 2554576 |
| D20 | 1972 | 1356975 |
| D21 | 1378 | 1963792 |
| D22 | 1348 | 2124265 |

In all strains, the proportion of IES and IES positions with a blast hit were relatively low because the IES are filled with repeat sequences and transposable elements (Hamilton et al., 2016). Nevertheless, despite its high similarity to the SB210 strain (only 48 SNPs in the MAC genome), the genome assembly of the strain D18 showed a number of IES positions lower than those observed in other strains by several orders of magnitude.

We believed that the above results indicated an absence of MIC genome in the assembly of the strain D18, with the few IES detected being either retained or differentially excised in the MAC genome of this strain.
