## Supplementary Material 11 Derelle et al. for "The macronuclear genomic landscape within *Tetrahymena thermophila*"

**Supplementary Material 11: heatmap of indels at MDS junctions as compared with SB210**

This heatmap is the same as in Figure 3B, including the three datasets published in (Wang et al., 2021):

_ SRR11922748: population B

_ SRR11922749: population A

_ SRR11922750: ancestral population

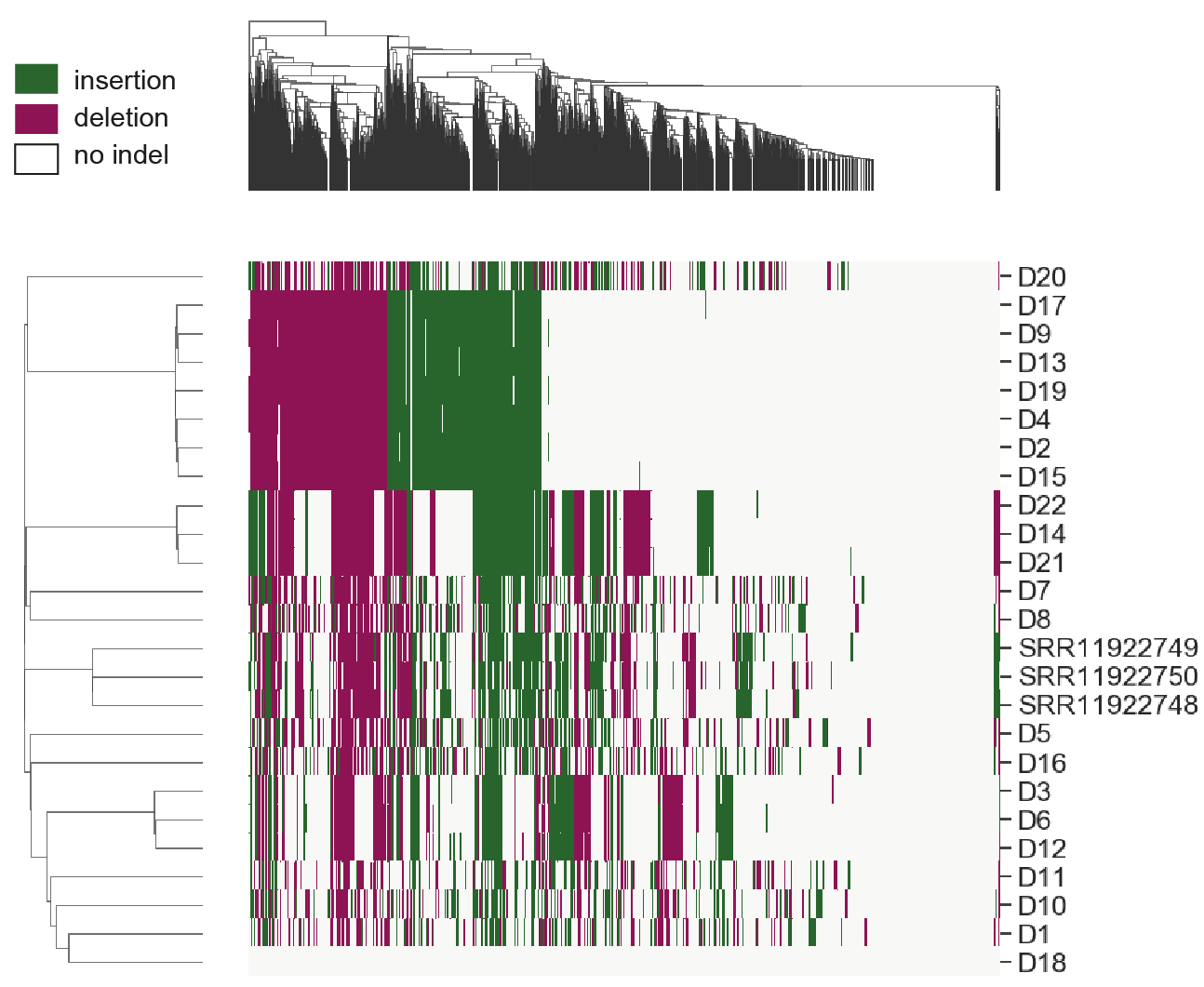

Wang, G., Fu, L., Xiong, J., Mochizuki, K., Fu, Y., and Miao, W. (2021). Identification and Characterization of Base-Substitution Mutations in the Macronuclear Genome of the Ciliate Tetrahymena thermophila. Genome Biol Evol 13.
