## Supplementary Material 12 Derelle et al. for "The macronuclear genomic landscape within *Tetrahymena thermophila*"

**Supplementary Material 12:
Single nucleotide diversity within strains**

A)

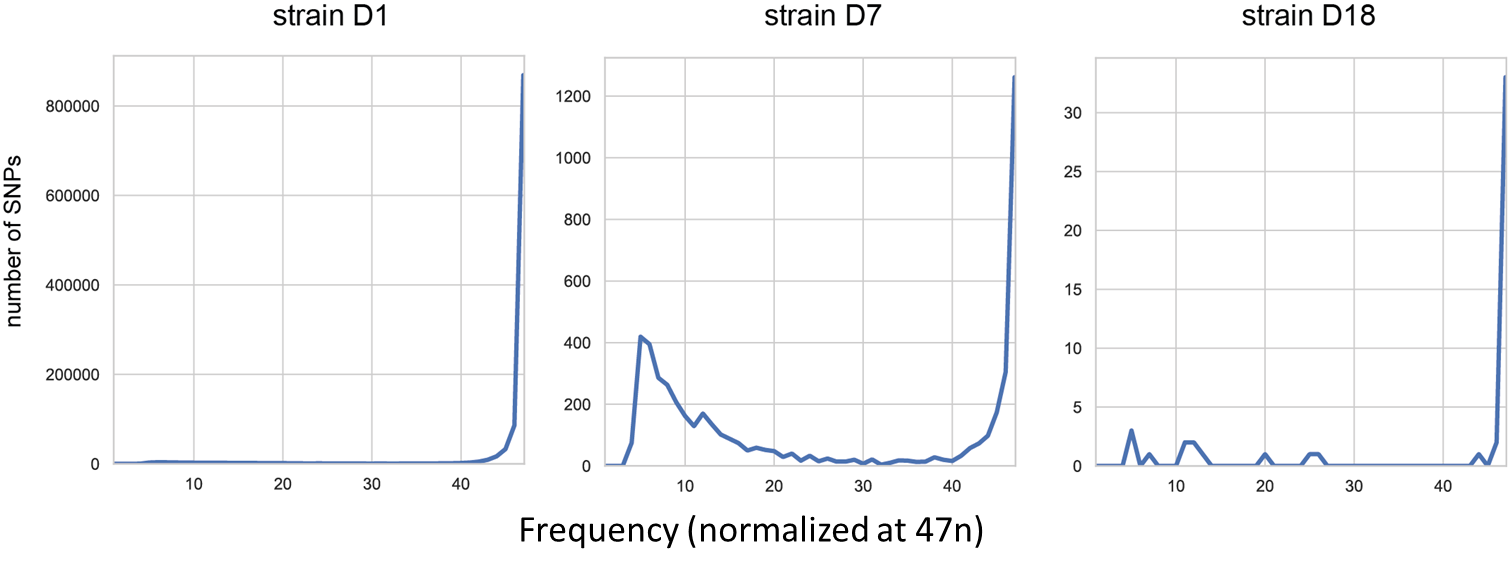

Number of variable positions as compared with SB210 as a function of their frequency in three strains over the 22 with decreasing divergence from the reference SB210 genome. D1 is highly distant, D7 is closely related, and D18 is almost identical to SB210.

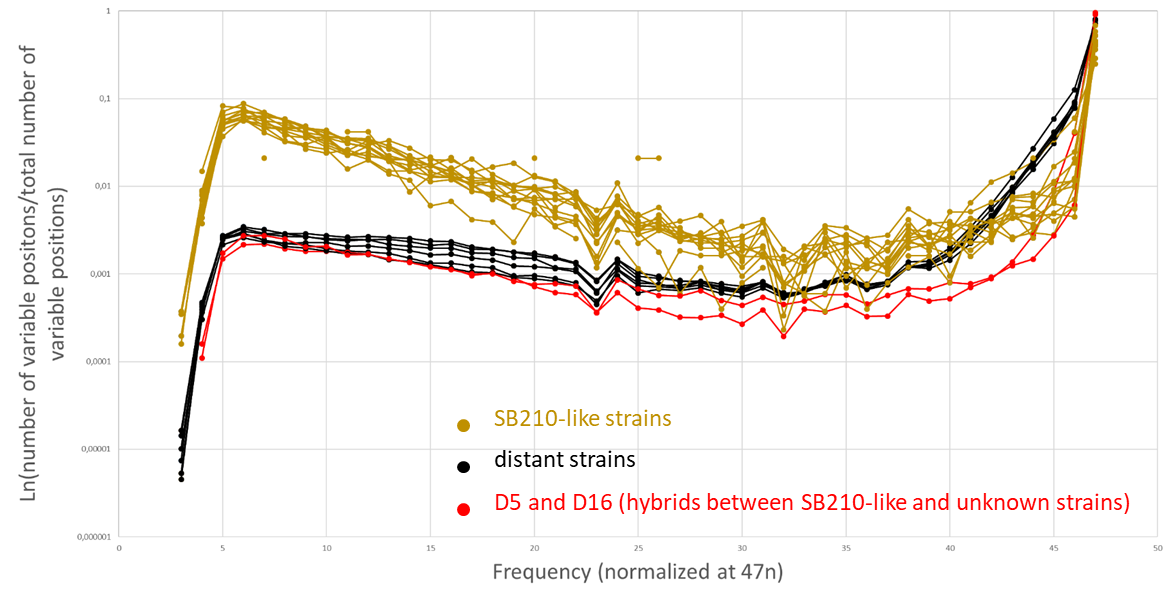
B)

Proportion of variable positions as compared with SB210 whose frequency ranges from 1 to 47n

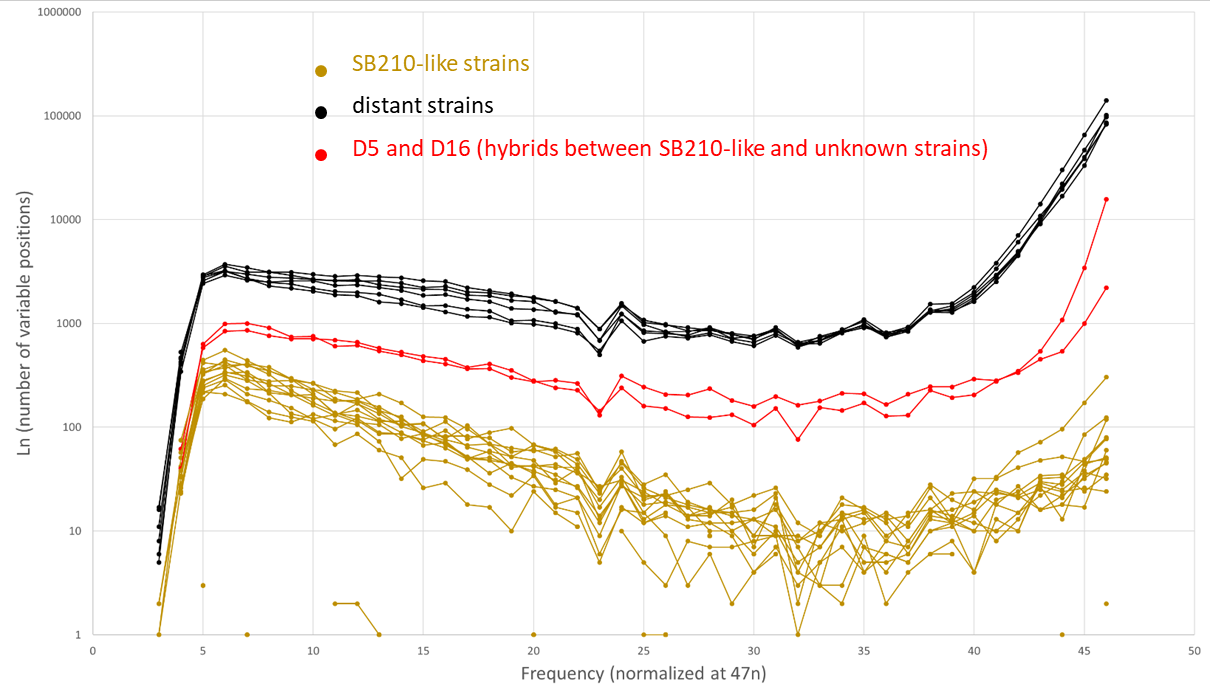
C)

Number of variable positions as compared with SB210 whose frequency ranges from 1 to 47n
