## Supplementary Material 13 Derelle et al. for "The macronuclear genomic landscape within *Tetrahymena thermophila*"

**Supplementary Material 13: pN/pS per GO term**

Distributions of pN/pS values for each slim Gene Ontology (GO). We only kept in the following boxplots GO with at least 10 genes (the number of genes assigned to each GO is indicated within brackets).

**Cellular component**

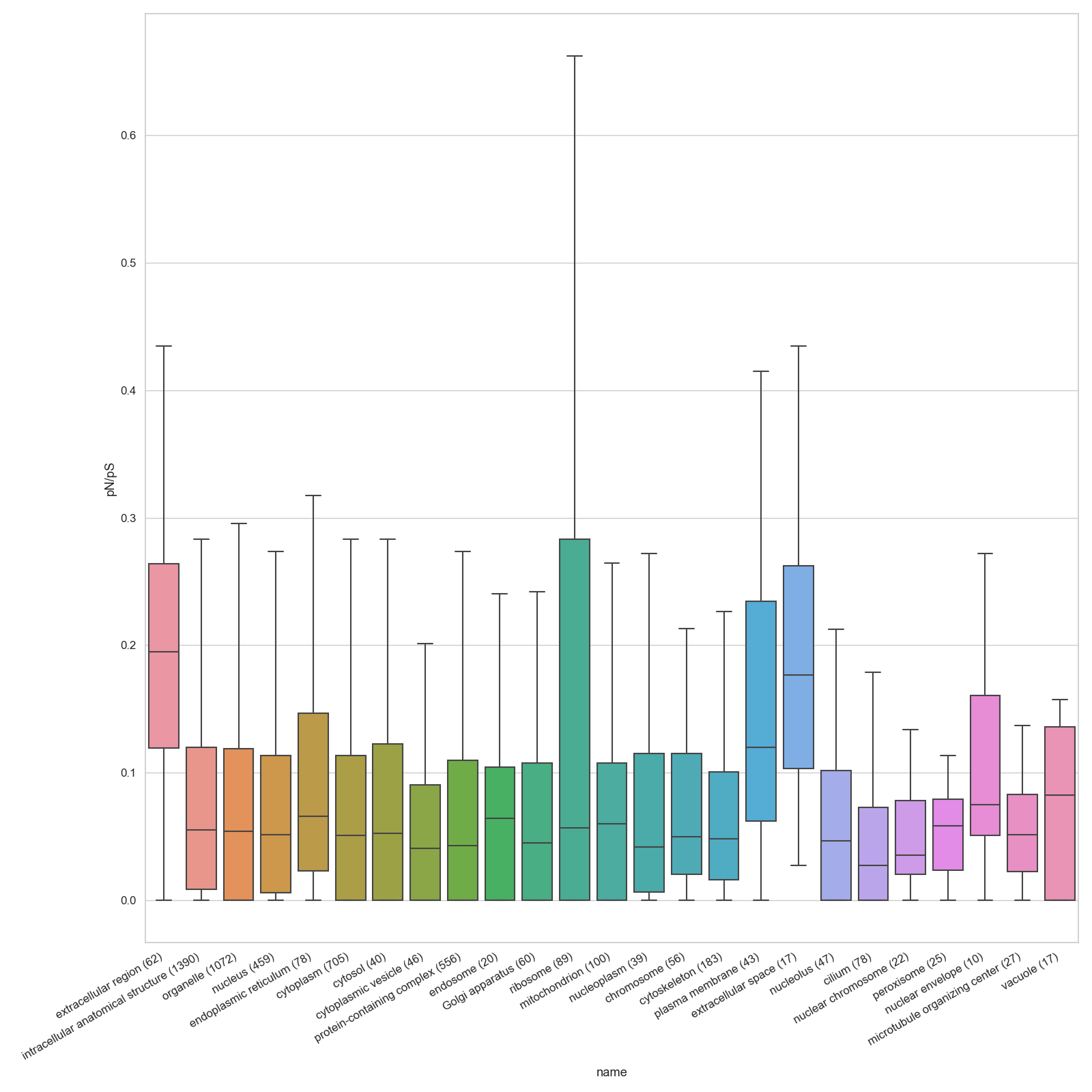

**Molecular function**

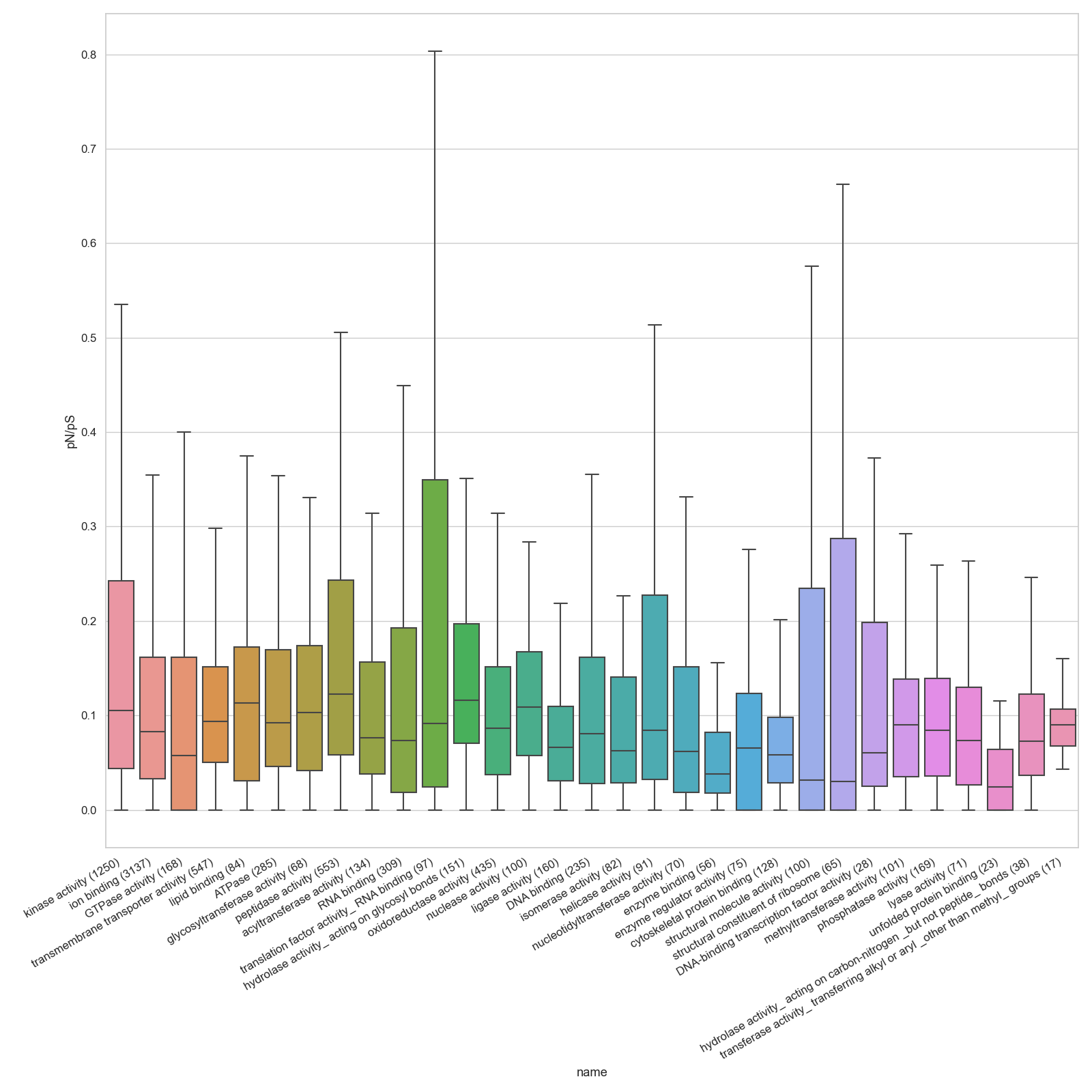

**Biological process**

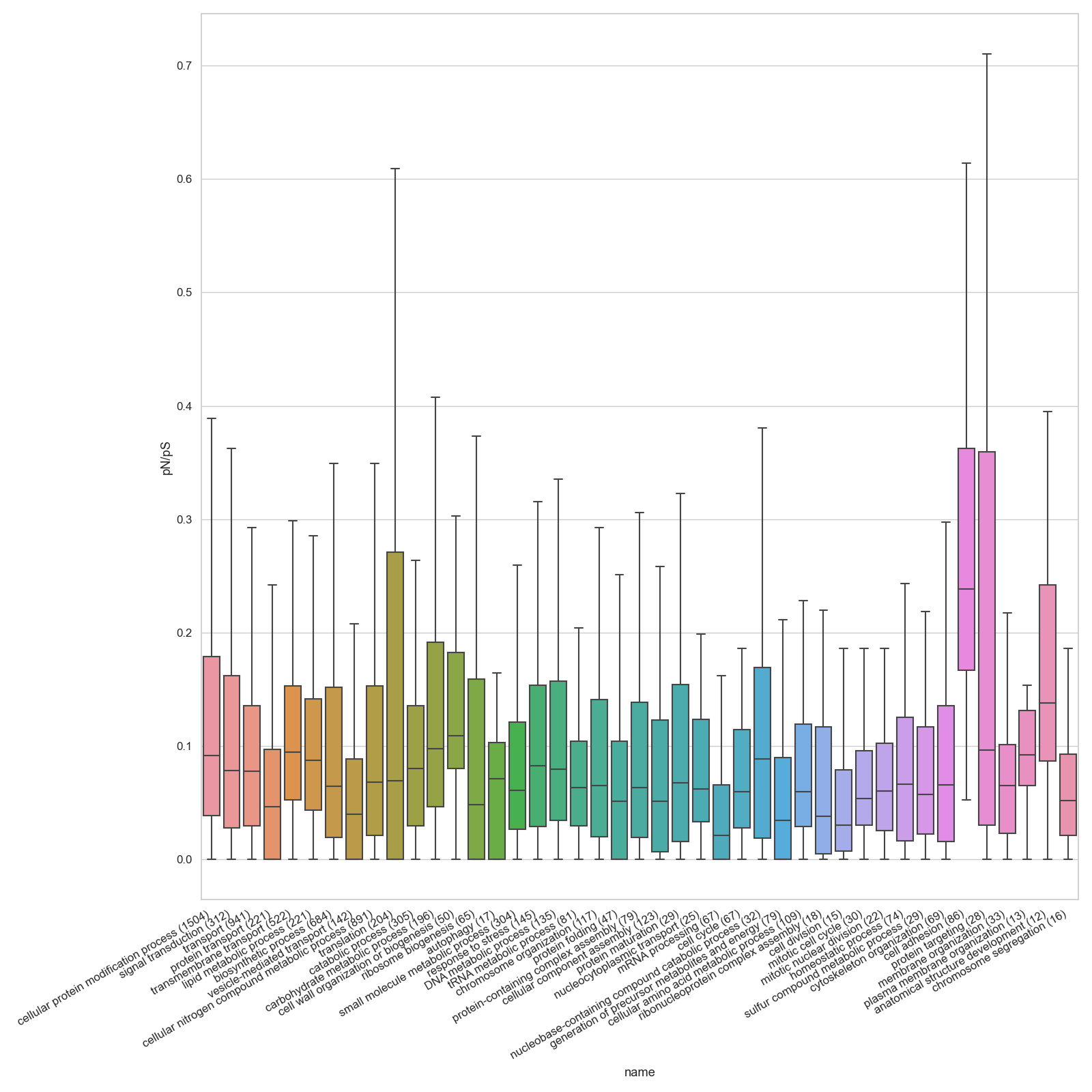
