## Supplementary Material 14 Derelle et al. for "The macronuclear genomic landscape within *Tetrahymena thermophila*"

**Supplementary Material 14: Direction of Selection per GO term**

Distributions of DoS values for each slim Gene Ontology (GO). We only kept in the following boxplots GO with at least 10 genes (the number of genes assigned to each GO is indicated within brackets).

**Cellular component**

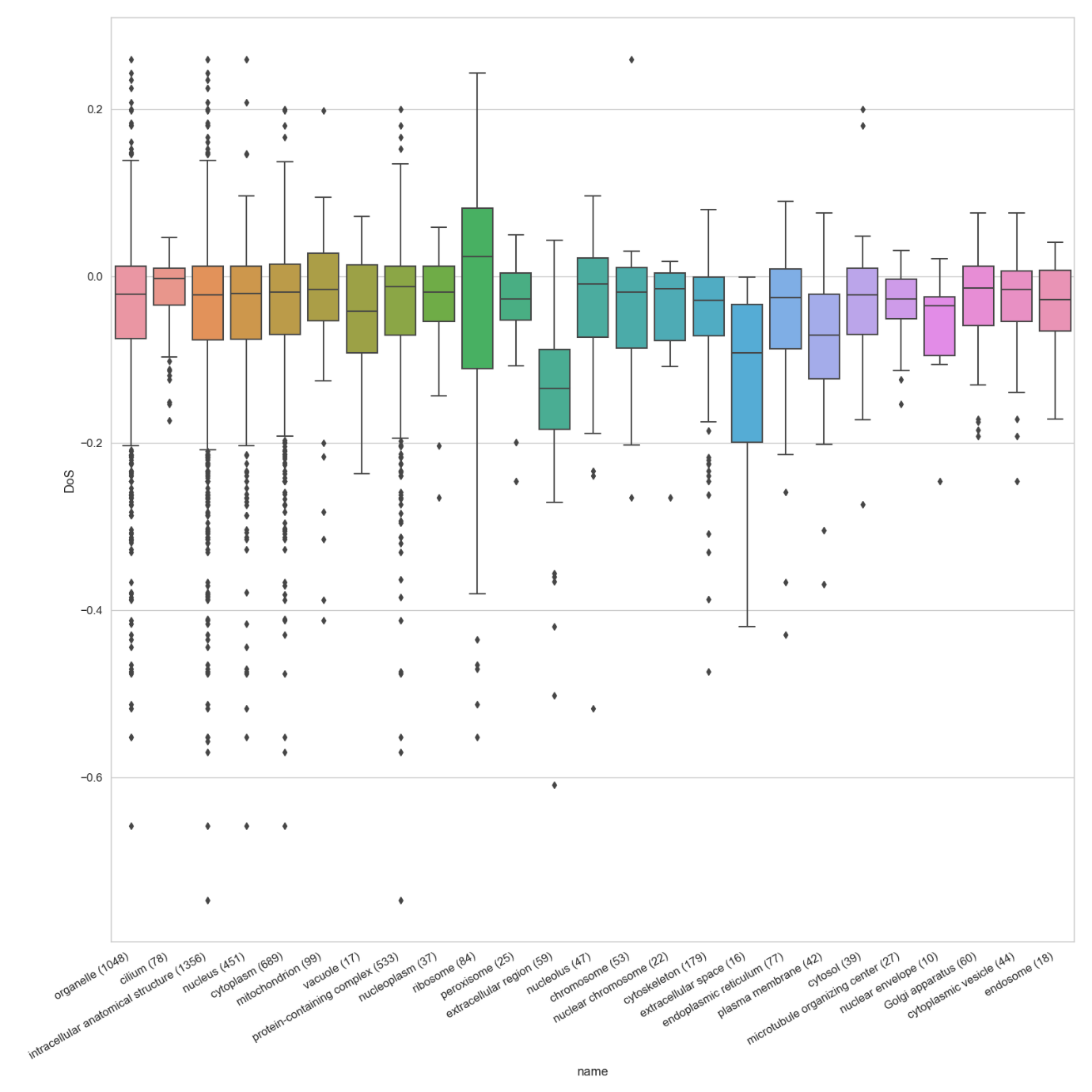

**Molecular function**

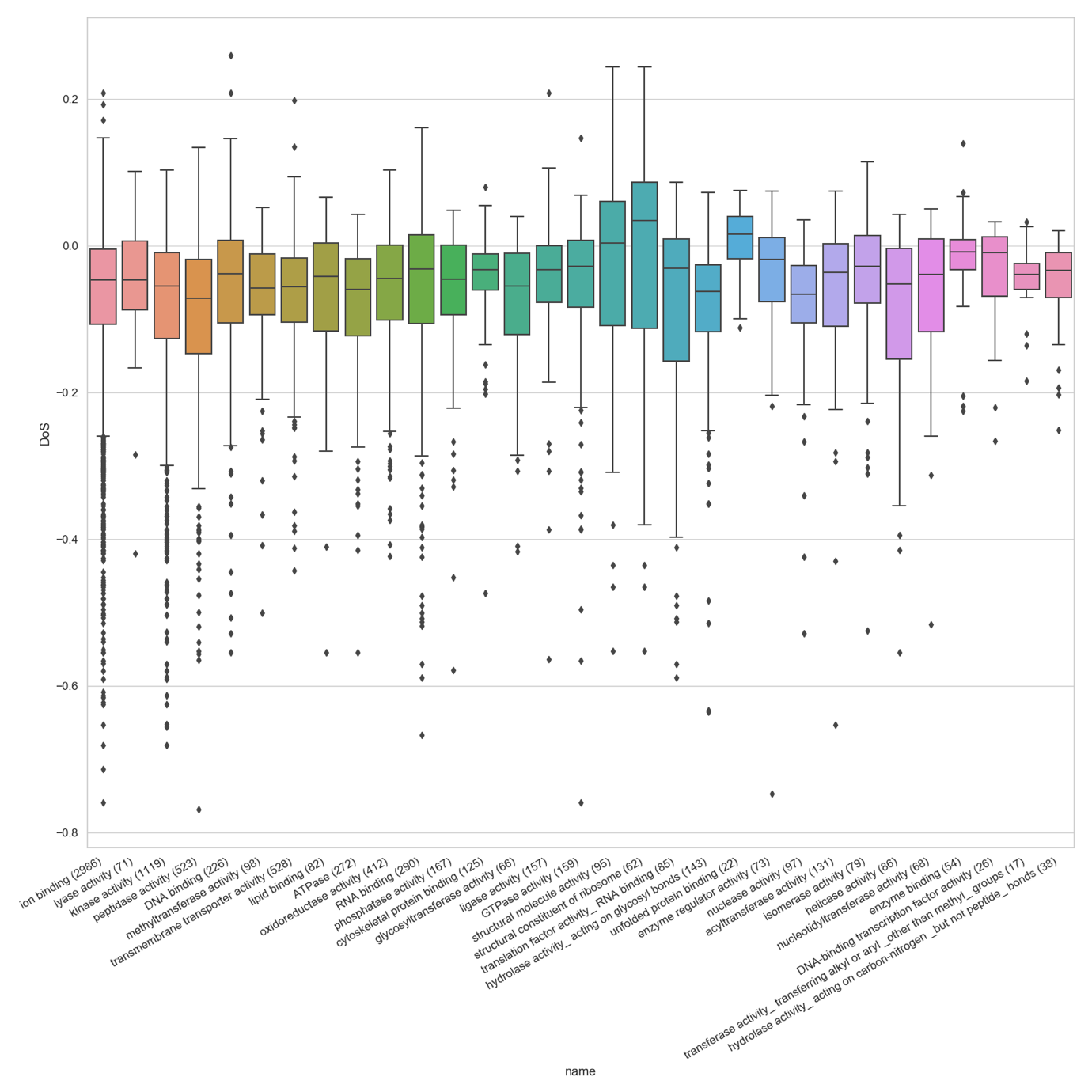

**Biological process**

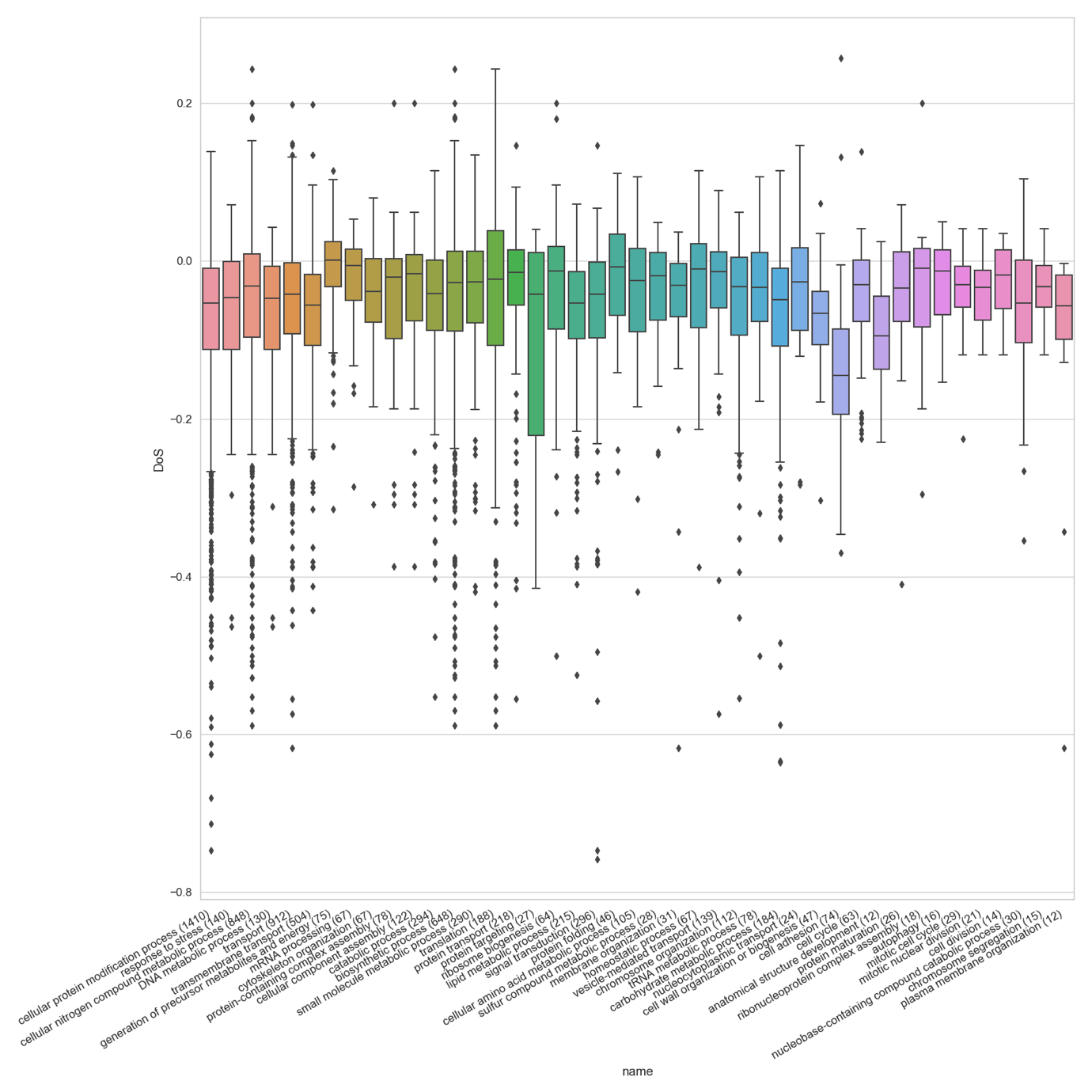
