## Supplementary Material 15 Derelle et al. for "The macronuclear genomic landscape within *Tetrahymena thermophila*"

**Supplementary Material 15: Direction of Selection around MIC centromeres**

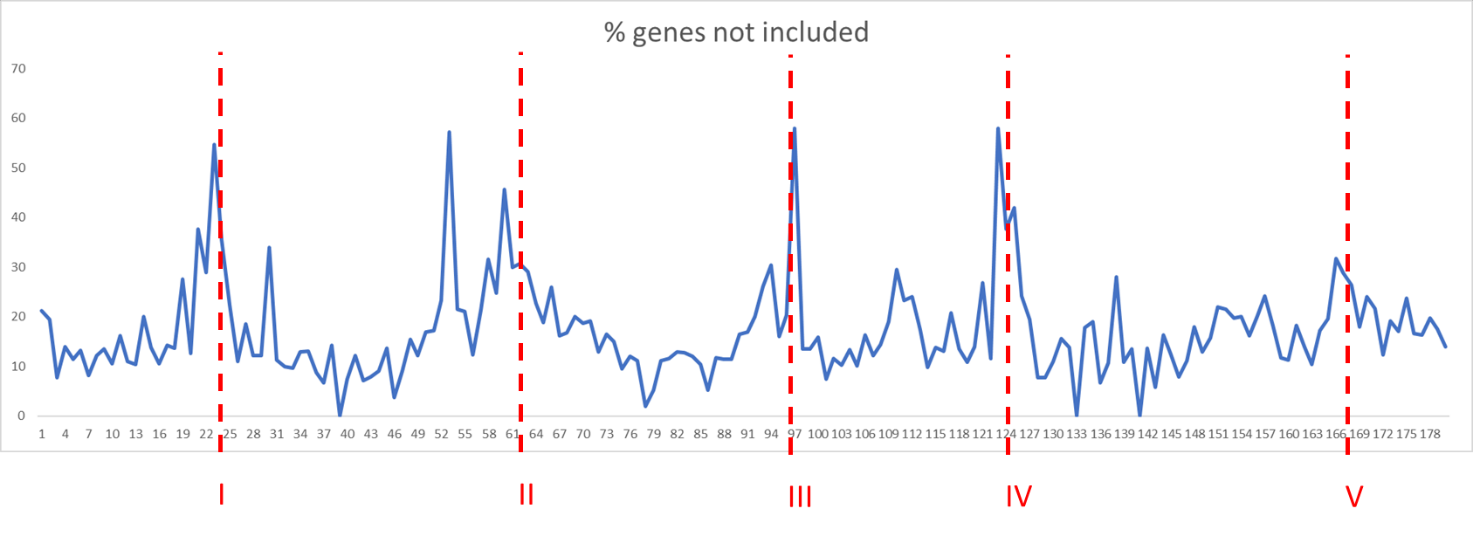

Number of genes not included in the DoS estimation because no orthologs were found in the genus *Tetrahymena*. Red dashed bars correspond to the positions of MIC centromeres.

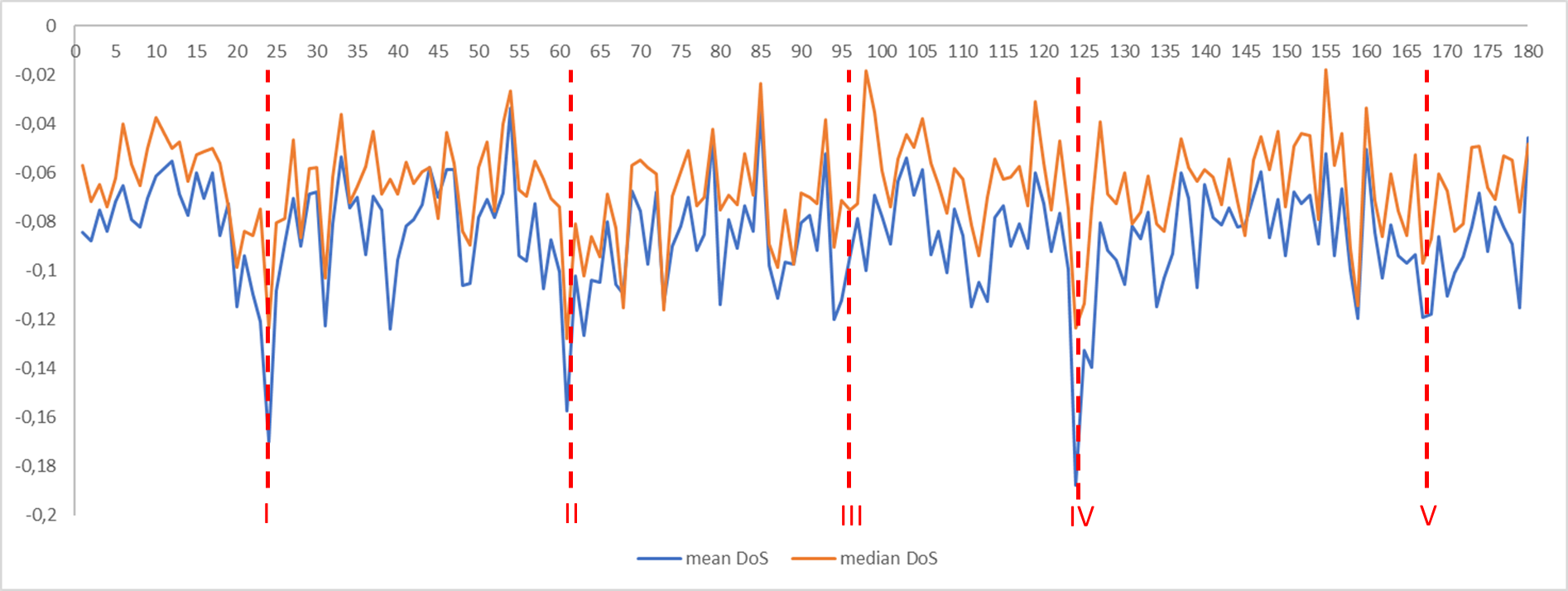

Mean (in blue) and median (in orange) of genes DoS per MAC chromosome. Red dashed bars correspond to the positions of MIC centromeres.
